# Area-specific synapse structure in branched axons reveals a subcellular level of complexity in thalamocortical networks

**DOI:** 10.1101/798926

**Authors:** Javier Rodriguez-Moreno, Cesar Porrero, Astrid Rollenhagen, Mario Rubio-Teves, Diana Casas-Torremocha, Lidia Alonso-Nanclares, Rachida Yakoubi, Andrea Santuy, Angel Merchan-Pérez, Javier DeFelipe, Joachim HR Lübke, Francisco Clasca

## Abstract

Thalamocortical Posterior nucleus (Po) axons innervating the somatosensory (S1) and motor (MC) vibrissal cortices are key links in the brain neuronal network that allows rodents to explore the environment whisking with their motile vibrissae. Here, using high-end 3D electron microscopy, we demonstrate massive differences between MC vs. S1 Po synapses in a) bouton and active zone size; b) neurotransmitter vesicle pool size; c) mitochondria distribution near synapses; and d) proportion of non-spinous dendrite contacts. These differences are as large, or bigger, than those between Po and ventroposterior thalamic nucleus synapses in S1. Moreover, using single-axon transfection labeling, we show that the structure of boutons in the MC vs. S1 branches of individual Po axons is different. These structural differences parallel striking, recently-discovered divergences in functional efficacy and plasticity between S1 and MC Po synapses, and overall reveal a new, subcellular level of thalamocortical circuit complexity, unaccounted for in current models.

## Introduction

Rodents explore their environment by rhythmically “whisking” with their motile facial vibrissae. Time-dependent correlations between motor commands and vibrissal follicle mechanoreceptor signals are used for inferring object position and texture. Such computations are carried out in multilevel, closed-loop neuronal networks encompassing the brainstem, thalamus and neocortex (reviewed in ^1^).

Two thalamocortical pathways lay at the core of these loops: Ventral Posteromedial thalamic nucleus (VPM) axons arborizing focally on L4 in the vibrissal “barrel” domain of the primary somatosensory cortex (S1); and Posterior thalamic nucleus (Po) axons arborizing mainly in L5a but also L1 of S1 ^2; 3^. Importantly, Po axons may target, in addition, the motor cortex (MC) middle layers ^3, 4, 5^ (L5-L3). The VPM axons relay time-locked mechanoreceptive single-whisker trigeminal signals, crucial for computing object location. In turn, Po axons convey information mainly about timing/amplitude differences between ongoing cortical outputs and multi-whisker sensory signals, which may be important for computing whisker position ^1, 6, 7^. Po cells are primarily driven by heavy and highly effective L5 cortico-thalamic inputs, while ascending trigeminal inputs to Po are sparser and largely modulatory in character ^7, 8^.

Activation of S1-L4 VPM synapses elicits large currents in cortical neurons ^9, 10^ and drives their firing with low failure rates ^11, 12^. VPM synapses are driven only by ionotropic receptors, depress rapidly upon repetitive stimulation ^9^ and, after an early postnatal period, show considerable resistance to sensory experience-dependent changes^13^. In contrast, synaptic potentials evoked in S1 by Po axons show slower rise and decay times, and elicit smaller currents ^10, 14^. The Po S1 synapses involve both ionotropic and metabotropic glutamate conductances, show paired-pulse facilitation ^10, 15^ and rapidly increase their efficacy in response to learned reward even in the adult ^13^. Recent data about the Po MC synapses indicate that, surprisingly, they elicit large currents, depress upon repetitive activation ^5, 16^ and involve only ionotropic receptors ^3^.

In cortical synapses, bouton and mitochondrial volume, synaptic vesicle pool size, and postsynaptic density (PSD) complexity and surface area directly correlate with increased neurotransmitter release probability and synaptic strength ^17, 18; 19^. Furthermore, the PSD surface area is proportional to the number of postsynaptic receptors ^20–23^. Thus, given the markedly divergent effect observed in S1 vs. MC Po synapses, and recent light microscope evidence that Po S1 axon varicosities are relatively small ^3^ we aimed to determine whether Po synapses in MC and S1 were different in their ultrastructural composition. Moreover, as studies in rat have reported that MC and S1 can be simultaneously targeted by branches of the same individual Po axons ^4, 24^, we also set out to elucidate if structural differences occur between boutons on separate branches of the same Po cell axon.

To visualize Po axons and measure identified Po synapses in S1 and MC, we combined single-cell and bulk axonal labeling with light and high-resolution, fine-scale 3D-electron microscopy (serial sectioning Transmission Electron Microscopy, ssTEM; and Focused Ion Beam Milling Scanning Electron Microscopy; FIB-SEM). In addition, as Po and VPM synapses have been recently shown to elicit marked different effects on S1 neurons, we re-analyzed our own dataset on VPM S1 synapses ^25^ and compared it with the Po synapse data.

Here, we demonstrate that axonal varicosities of Po axons in MC are much larger than those of the same axons in S1, and that their presynaptic structure and postsynaptic relationships in each cortical area are strikingly different. In addition, we report similarly sharp differences between Po and VPM axon synapses in S1. Along with recent parallel functional observations ^3, 16^ differences in synapse structure both between axons originated in diverse thalamic nuclei, and between the branches of the same individual axons targeting separate cortical areas/layers, call into question current models of thalamocortical interaction.

## Results

### Light microscopic visualization of Po axon terminals in S1 and in MC

To selectively label a sizable population of thalamocortical Po axons, 10 kDa lysine-fixable biotinylated dextranamine (BDA) was iontophoretically delivered into Po. Only experiments in which the BDA deposit was restricted to the Po nucleus were analyzed (Fig. 1a-b; Supplementary Fig. SM1). In S1, two distinct bands of axonal arborizations were labeled: one in L5a and the other in upper half of L1 (Fig. 1c). In MC, labeled Po axonal arborizations formed a single band from upper L5a to lower L3 (Fig. 1d). The Golgi-like axon staining revealed frequent varicosities of variable size (Fig. 1e-g).

**Fig. 1.**
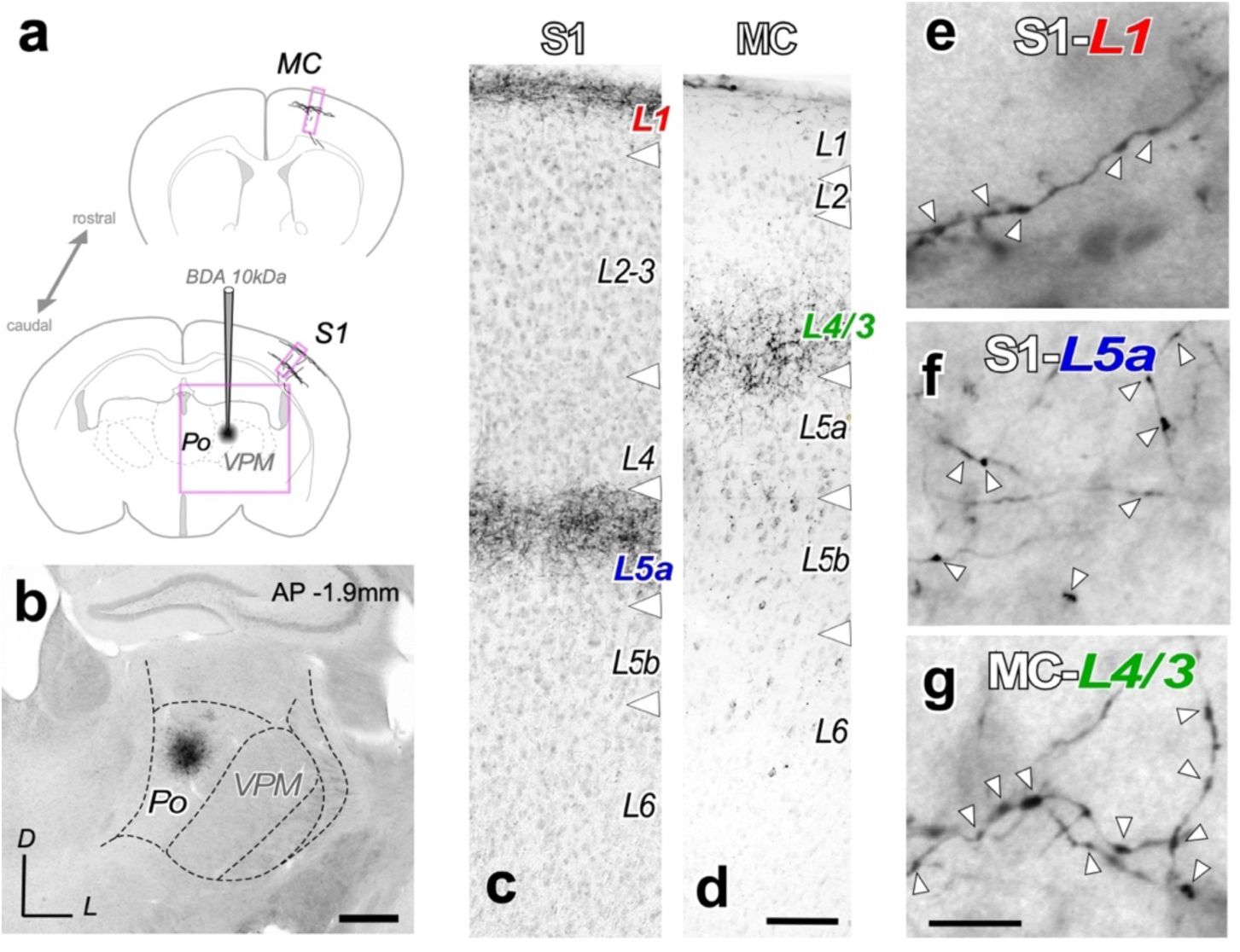
Selective bulk-labeling of Po thalamocortical axons in the vibrissal motor (MC) and vibrissal somatosensory (S1) cortices. (**a**) BDA iontophoresis procedure. Schematic coronal brain sections depicting the BDA deposit in Po, and the anterogradely labeled thalamocortical axonal in S1 and MC. (**b**) A typical iontophoretic BDA Po deposit. The region illustrated is the area framed in panel “a”. Thionin counterstain. Scale bar represents 250 µm. (**c-d**) Coronal sections showing the layer selective arborization of BDA-labeled Po axons in specific layers (L), namely L1 and L5a in S1 (“c”) and L4/3 in MC (“d”). Scale bar represents 100 µm. (**e**-**g**) High-magnification images of BDA-labeled Po axons in S1-L1. (**e**), in S1-L5a (**f**), and in MC-L4/3 (**g**). Axonal varicosities are marked by arrowheads. Scale bar represents 10 µm.

### ssTEM and FIB-SEM analysis of labeled thalamocortical Po axons

Electron microscopic samples were taken from sections adjacent to those containing the heaviest anterograde labeling in S1 or MC (see Methods; Supplementary Fig. SM2). These were subsequently analyzed using either high-resolution ssTEM or FIB-SEM. In S1, samples were taken from L1 and L5a, which were readily delineated cytoarchitecturally. However, the cytoarchitectonic definition of MC middle layers is ambiguous; thus, our MC samples included L4 and adjacent deep parts of L3; we refer to this neuropil as “MC-L4/3”.

A total of 220 Po synaptic boutons and their respective target structures were reconstructed and analyzed with ssTEM or FIB-SEM: 80 in S1-L5a, 71 in S1-L1 and 69 in MC-L4/3. The two serial 3D EM techniques yielded in consistent results. The ssTEM analysis allowed a high-resolution 3D-volume reconstruction of the overall geometry of synaptic complexes including the subsequent quantification of PSDs surface area and vesicle pool size, however, it sampled only axon varicosities (putative boutons), and a smaller neuropil volume, which not always allowed a full visualization of the spine neck. In turn, FIB-SEM analysis allowed the complete visualization of postsynaptic elements and inter-bouton axonal segments. For efficiency, the FIB-SEM image acquisition was made with a slightly lower resolution than the ssTEM images; for this reason, fine details such as the individual synaptic vesicles could be only unambiguously resolved and counted in the ssTEM samples.

### Ultrastructural features of Po synaptic boutons in the vibrissal regions of S1 and MC

A total of 192 axon boutons, most, but not all of them containing at least one synaptic site, were reconstructed and analyzed with ssTEM or FIB-SEM: 74 BDA-labeled boutons from S1-L5a, 67 boutons from S1-L1 and 51 boutons from MC-L4/3 (Table 1). Labeled boutons were identified by their electron-dense DAB reaction product in their cytoplasm and by their tapering into the adjacent axonal segments. A second criterion was the presence of a postsynaptic density (PSD) in consecutive serial sections indicative for a synaptic contact. The sampling of labeled boutons in ssTEM was essentially random, as any labeled bouton that could be followed from its beginning to its end within a given series of ultrathin sections was reconstructed and quantified. In FIB-SEM image stacks, all the labeled axonal segments, varicose or not, contained in the stack volume were analyzed.

**Table 1:**
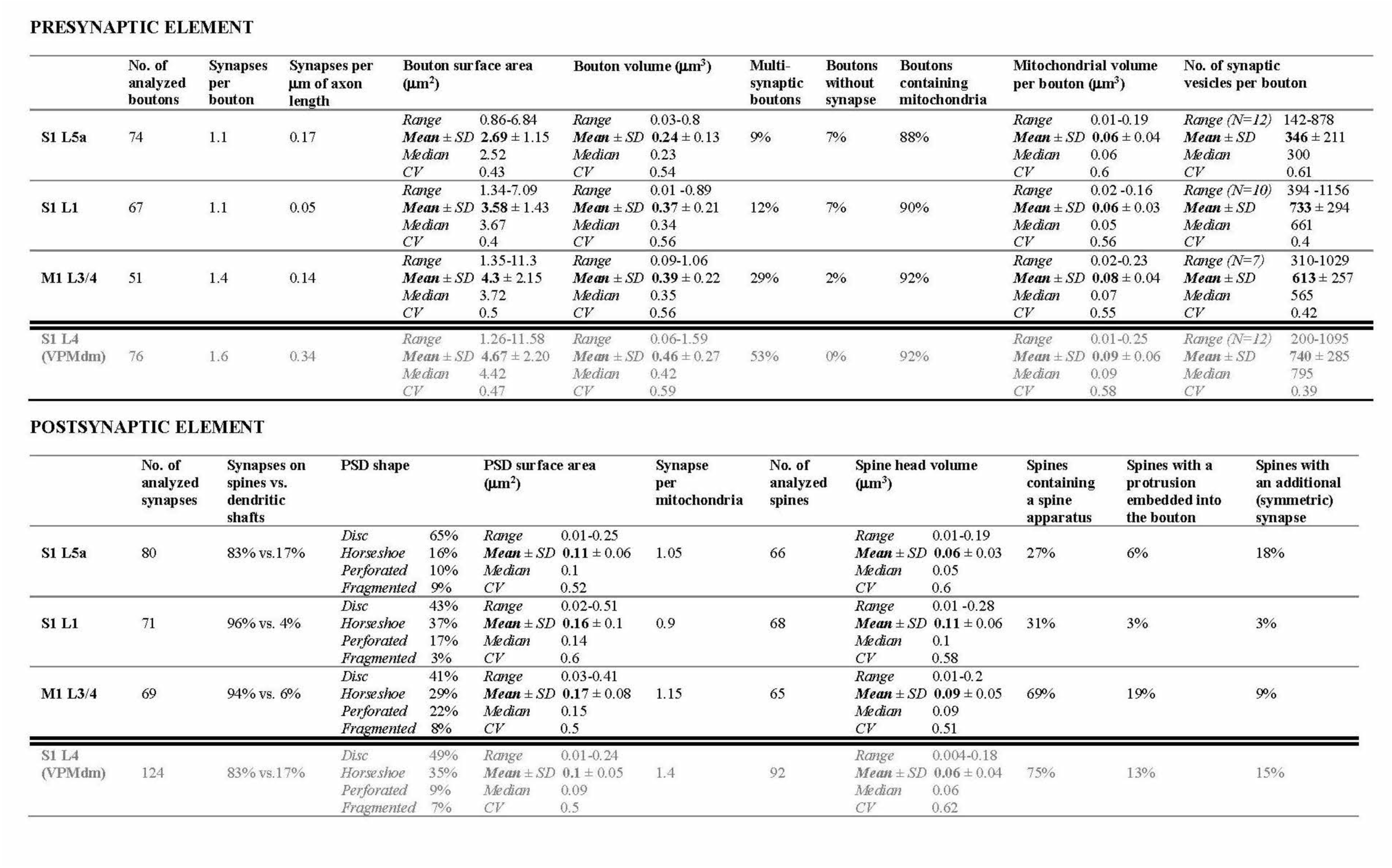
Ultrastructural 3D measurements of Po synapses in MC-L4/3, S1-L1 or S1-L5a, and comparison with VPM synapses in S1-L4.

Synaptic contacts between a labeled Po axon and its postsynaptic target were identified by the presence of distinct, parallel pre- and postsynaptic membranes at the synaptic apposition zone, separated by a synaptic cleft and an electron-dense band adherent to the cytoplasmic surface of the postsynaptic membrane (PSD). This corresponds to asymmetric synapses, regarded as excitatory and thus glutamatergic ^26^. Mitochondria and synaptic vesicles were clearly visible and, in the boutons less heavily-stained, it was even possible to identify and count with confidence all the synaptic vesicles.

In the three cortex regions studied, the large majority of Po boutons were monosynaptic. About 30% of boutons in MC-L4/3 and ∼10% in S1 (L5a and L1) simultaneously innervated two target structures (Fig. 2d, 2f, Table 1). We did not observe any Po bouton with three or more synapses, which is a frequent finding in S1-L4 VPM boutons ^25^ (Table 1).

**Fig. 2.**
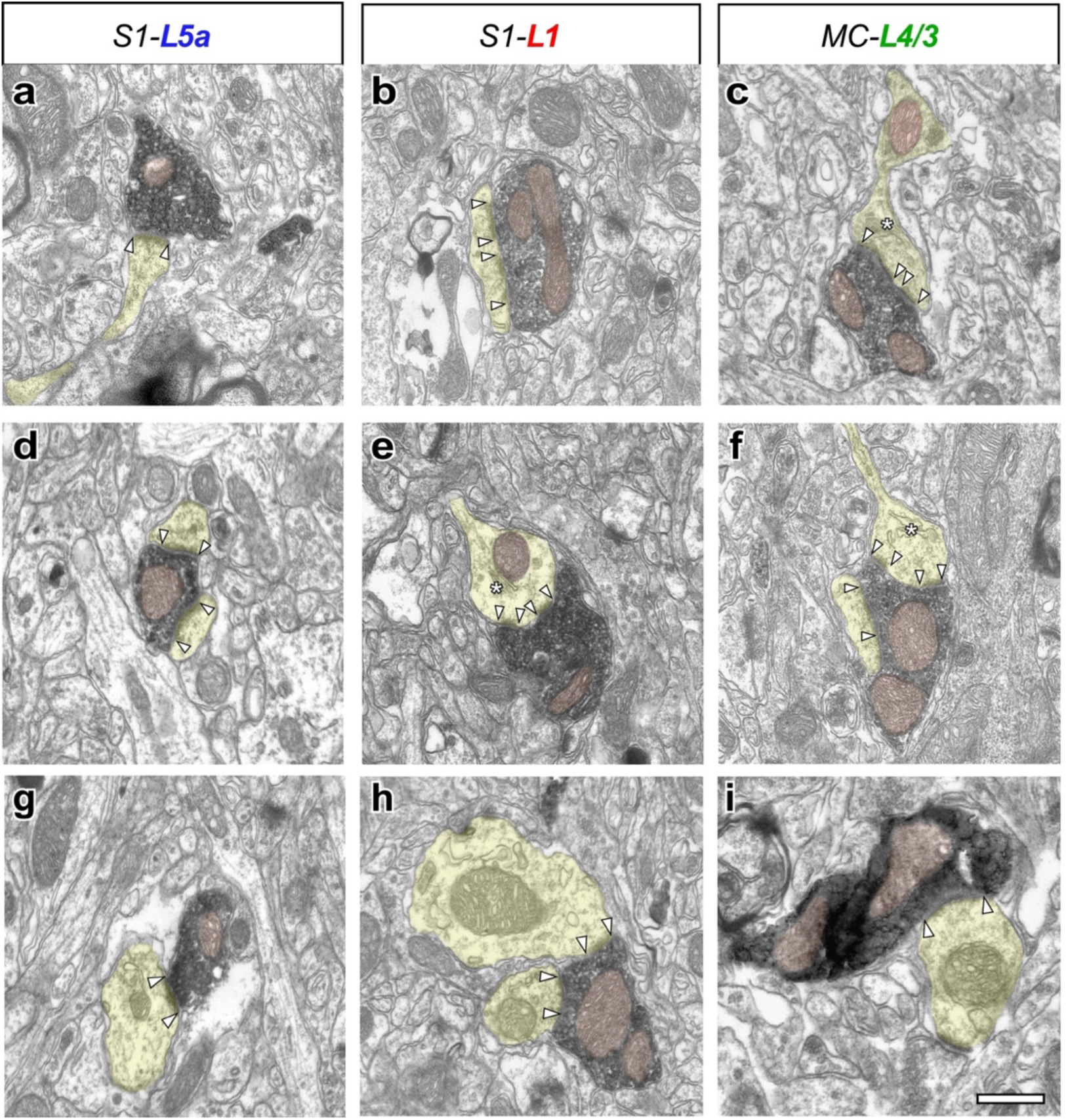
Electron micrograph examples of BDA-labeled Po thalamocortical boutons establishing synaptic contacts with cortical dendritic spines or shafts. In all images, dendritic shafts and spines are highlighted in transparent yellow and mitochondria are shaded in transparent brown. PSD borders are marked by white arrowheads. Asterisks indicate the spine apparatus. Inside some boutons, synaptic vesicles are clearly visible. (**a**, **d**) Typical Po boutons synapsing on a dendritic spine head (**a**) or simultaneously onto two different spine heads (**d**) in S1-L5a. (**b, e**) Po boutons synapsing on dendritic spines in S1-L1. In **e**, note the presence of a mitochondrion and a spine apparatus (asterisk) within the postsynaptic spine head. (**c, f**) Synaptic Po boutons in MC-L3/4 synapsing onto one (**c**) or two dendritic spines simultaneously (f). (**g-i**) Examples of Po bouton contacts onto dendritic shafts, one in each of the three studied neuropils. Scale bar represents 0.5 µm.

Most Po boutons (88–92%, Table 1) contained one or several mitochondria of different shape and size (Fig. 2 and 3). Mitochondria contributed substantially to the bouton volume (25% in S1-L5a, 16% in S1-L1 and 20.5% in MC-L4/3). Remarkably, in both S1-L1 and S1-L5a, a significant number (7%) of the Po axonal varicosities that contained a mitochondrion and synaptic vesicles, lacked any evident synaptic contact. Such “non-synaptic boutons” were also observed in MC-L4/3 Po axons but less frequent (2%), but never in the VPM S1-L4 axons (Table 1; see also ^27^).

**Fig. 3.**
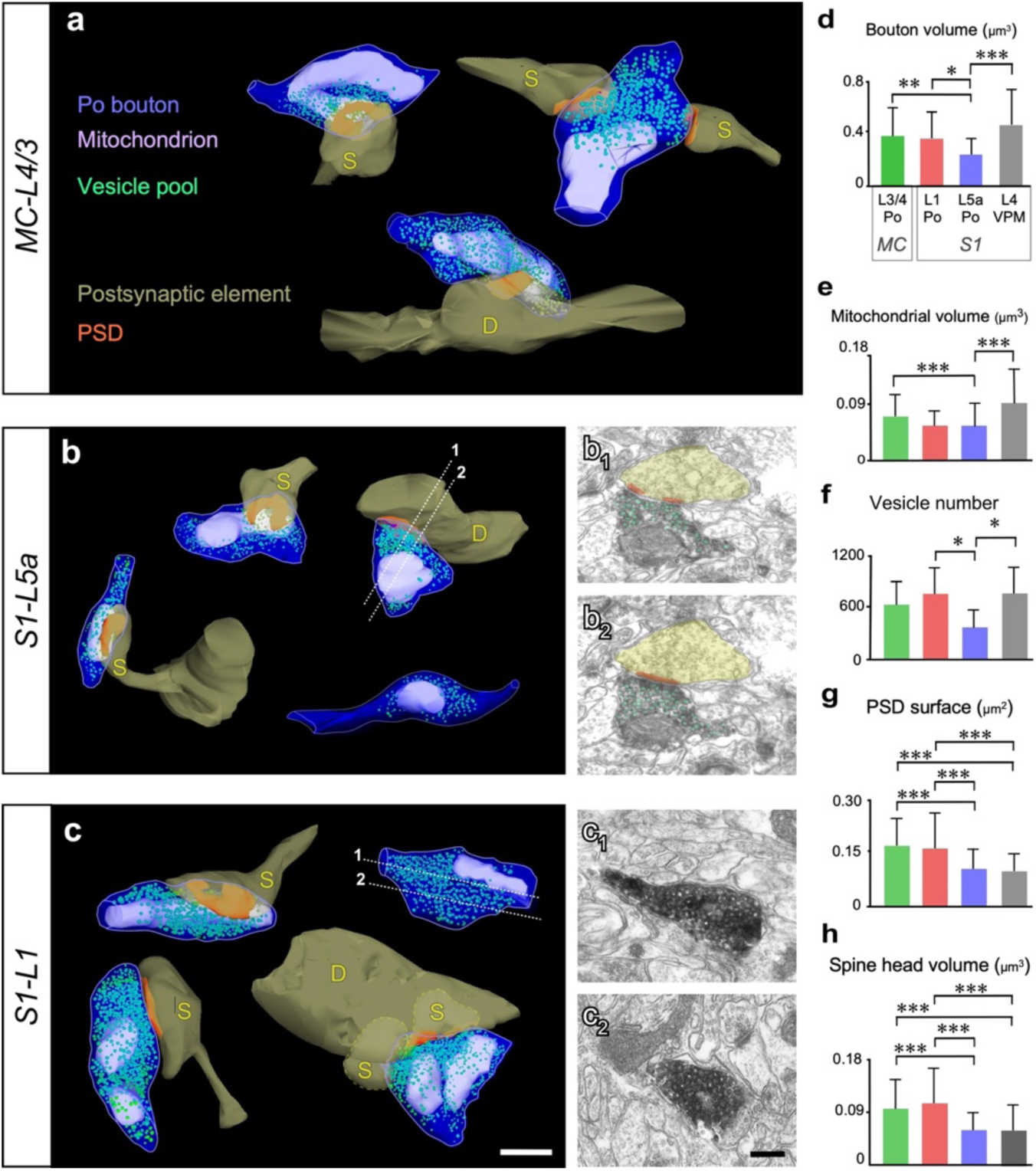
ssTEM 3D-reconstructions of thalamocortical Po boutons, and comparison of their ultrastructural parameters in the various cortical domains studied. (**a**) Boutons labeled in MC-L4/3. Top: Two boutons contacting spines (“S”). The bouton on the right is simultaneously establishing synaptic contacts with two different spines. Bottom: a synaptic bouton terminating on a dendrite shaft (“D”). (**b**) Boutons labeled in S1-L5a. Left: two boutons contacting dendritic spines. Right (top): a bouton synapsing on a dendritic shaft (“D”). Two of the serial sections (indicated by dashed white lines) out of which this bouton was reconstructed are illustrated in panels **b_1_**-**b_2_**. Right (bottom): an axon varicosity containing a mitochondrion and synaptic vesicles, but no adjacent PSD or postsynaptic profile. (**c**) Boutons labeled in S1-L1. Left: two boutons establishing synaptic contacts with mushroom-like dendritic spines. Right (top): axonal varicosity containing a mitochondrion and synaptic vesicles, but no adjacent PSD. Two sections are shown in (**c_1_**-**c_2_**). Right (bottom): a bouton simultaneously contacting a dendritic shaft and two different spines. Scale bar represents 0.5 µm. (**d**-**h)** Bar histograms comparing (**d**) bouton volume; (**e**) mitochondrial volume; (**f**) synaptic vesicle number; (**g**) PSD surface area; and (**h**) spine head volume between Po synapses in the three cortical neuropil areas investigated (colored bars). Measurements for VPM synaptic boutons in S1-L4 (gray bar) are included (gray bar). Levels of significance are indicated by asterisks: * p ≤ 0.05; ** p ≤ 0.01; *** p ≤ 0.001.

### Synaptic structural features specifically revealed by FIB-SEM analysis

Our FIB-SEM analysis is based on sampling 2,435 µm^3^ of cortical neuropil. At high magnification FIB-SEM, the axonal BDA axonal segments were few and widely scattered. Thus, despite taking our FIB-SEM samples from zones that under light microscopy appeared heavily labeled (Fig. 1), many of the volumes examined with FIB-SEM actually contained inside few or no labeled Po axonal segments. Nevertheless, twenty-six Po axonal segments totaling ∼283 µm of axonal length were reconstructed. Of these, 108 µm were measured from axonal branches in S1-L5a (0.16 synapses/µm of total axon length); 74 µm from S1-L1 (0.05 synapses/µm of total axon length); and 101 µm from MC-L4/3 (0.14 synapses/µm of total axon length; Table 1, Fig. 4, Supplementary Fig. SM3). Varicosities were defined as any swelling on the axon exceeding by more than 50% the typical variation of the adjacent axonal segments ^28^. About a quarter of all Po synapses analyzed with FIB-SEM (9 out of 37) occur in non-varicose “inter-bouton” axonal zones (S1-L5a: 5/20; S1-L1:1/4; MC-L4/3: 3/13). Several such synapses are indicated by arrows in Fig. 4a-c. In VPM S1-L4 axons, synapses in non-varicose regions are much less frequent ^25, 27^.

**Fig. 4.**
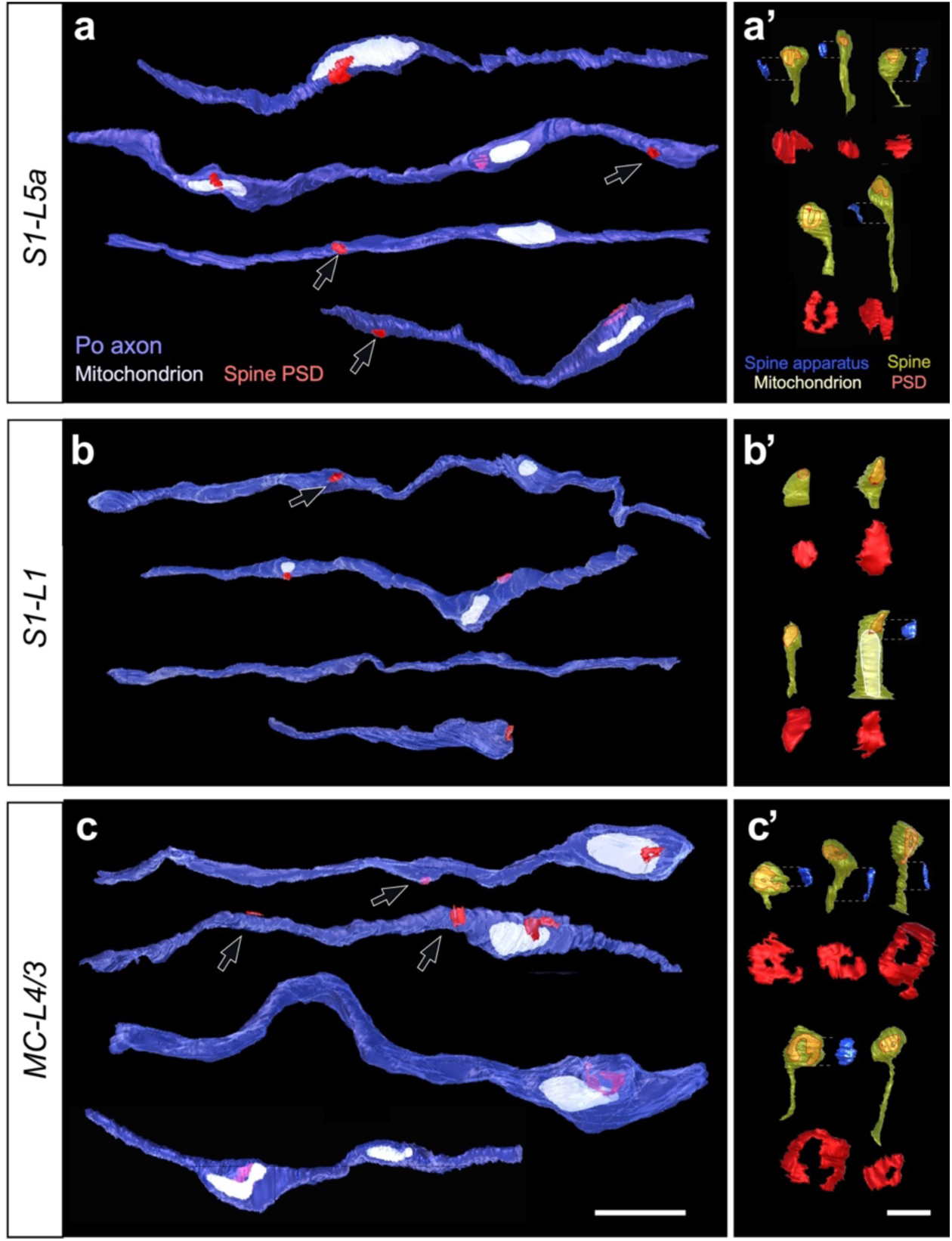
Representative examples of thalamocortical Po axon segments, postsynaptic dendritic spines and PSDs 3D-reconstructed from FIB-SEM image stacks. (**a**-**c**) Individual axonal segments from (**a**) S1-L5a; (**b**) S1-L1, or (**c**) MC-L4/3. Black arrows indicate PSDs located in non-varicose axon domains. Scale bar represents 2 µm. (**a’**-**c’**) Different morphologies of dendritic spines postsynaptic to Po boutons in (**a’**) S1-L5a, (**b’**) S1-L1, and (**c’**) MC-L4/3. Spine surface is partly transparent to allow the visualization of the PSDs within the spine head. In the spines containing a spine apparatus, this organelle (blue) is shown aside. Scale bar (for spines and spine apparatuses) represent 0.5 µm. For clarity, the spine PSDs are also represented isolated below, at double magnification. Note the differences in both shape (perforated vs. non-perforated, closed ring- or horseshoe-like) and size of the PSDs. In (**b’**) note the large spine containing a mitochondrion.

### Ultrastructural features of elements postsynaptic to the Po axons

In all three cortical regions investigated, the majority of Po synapses were established on dendritic spines (83–96%). Synapses onto dendritic shafts (which may partially correspond to non-spinous cortical interneurons ^29^) were frequent (17%) in S1-L5a, but less frequent by ∼4-fold in S1-L1 or MC-L4/3 (4–6%, respectively; Table 1; Fig. 2-4) with no contacts on neuronal somata.

Strikingly, the PSDs displayed a wide range in both shape and size (Fig. 4a’-c’, Table 1). The mean surface area of the S1-L5a PSDs was comparable (0.11 µm^2^) to that previously measured in VPM-L4 synapses (Table 1, Fig. 3g). Surprisingly, PSDs surface area of MC-L4/3 and S1-L1 Po synapses were ∼60% larger. In S1-L5a, most PSDs (65%) had disc-like morphologies. In contrast, most PSDs in S1-L1 (57%) or MC-L4/3 (59%), displayed more complex horseshoe-shaped, perforated or fragmented morphologies (Fig. 4a’-c’).

A total of 199 dendritic spines postsynaptic to Po boutons were analyzed (Table 1). Interestingly, some of them (3%-18%) were also targeted by an unlabeled symmetric (putatively inhibitory) synapse of unknown origin (Table 1). Only 30% of the dendritic spines postsynaptic to Po boutons in S1 (L5a and L1) contained a spine apparatus (an endoplasmic reticulum derivate, Fig. 4a’-4b’), while the large majority (70%) of postsynaptic spines in MC-L4/3 displayed this structural subelement responsible for spine motility but also stabilization of the pre-and postsynaptic apposition zone during synaptic transmission. Mitochondria were found inside 2 of the 68 dendritic spines postsynaptic to S1-L1 Po boutons, despite the exceedingly rare occurrence of spine mitochondria in rodent somatosensory cortex ^30^.

### Dendritic spine invaginations into thalamocortical Po boutons

Numerous dendritic spine heads postsynaptic to Po boutons formed a thick finger-like protrusion that invaginated into the presynaptic bouton (Fig. 5a-d). Such invaginations were observed in all three cortical domains investigated, but were more frequent in MC (Fig. 5e and Table 1). Invaginations were always adjacent to the spine PSD and present in spines with widely different PSDs sizes (Fig. 5e). Their volume was similar in the three cortical domains analyzed (range 0.6-0.75 µm^2^). The invaginated membranes were smooth, lacking evident membrane specializations (Fig. 5a_1_). We had observed similar invaginations in mouse S1-L4 VPM synapses; remarkably their prevalence in Po MC synapses seems to be higher ^25^ (13% vs. 19%; Fig. 5e-f).

**Fig. 5.**
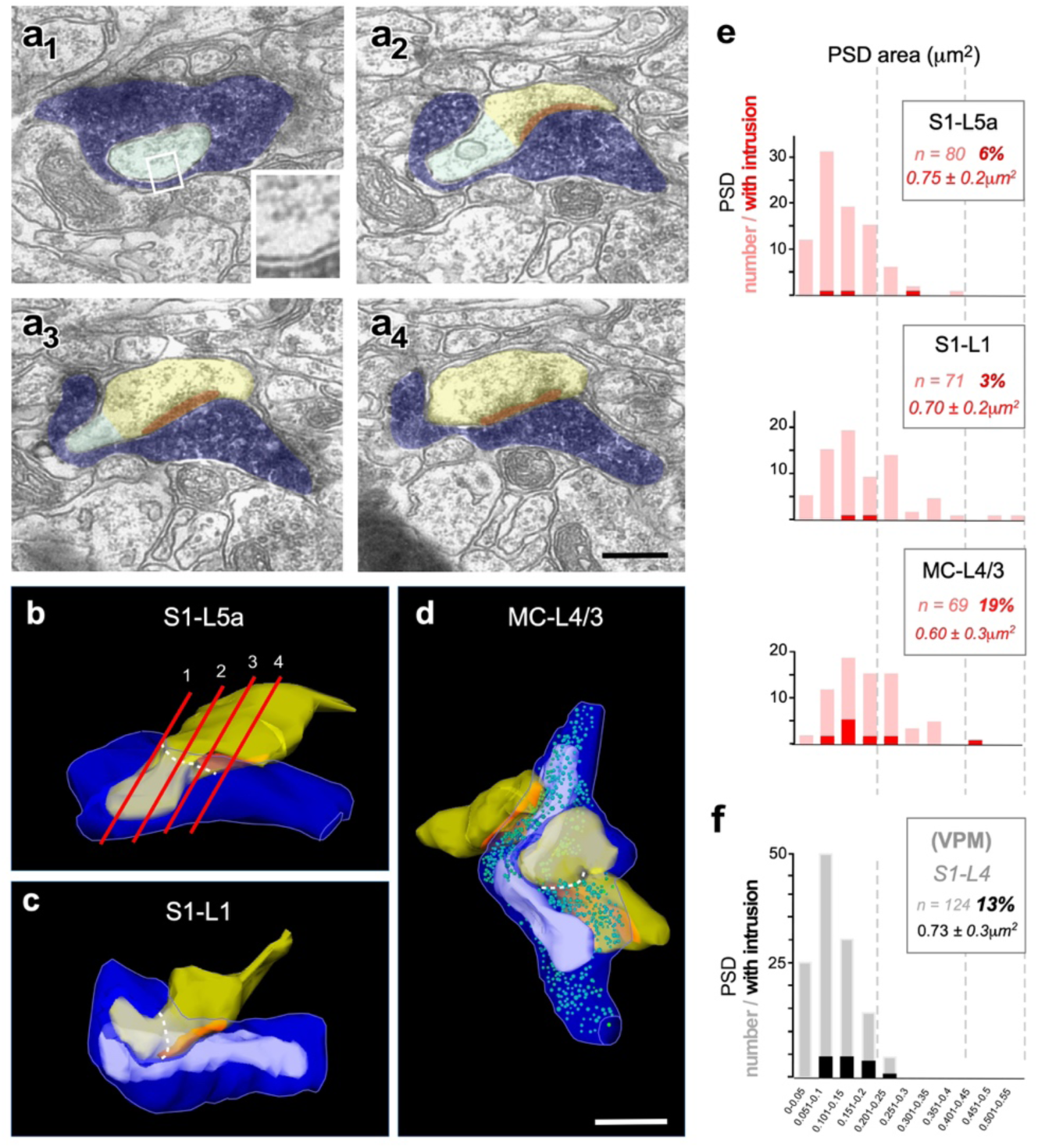
Large cortical dendritic spine protrusions invaginate into the presynaptic Po boutons. (**a_1_**-**a_4_**) Consecutive electron micrographs showing a postsynaptic spine head (transparent yellow) with a large protrusion (transparent green) invaginated into a labeled Po bouton (transparent blue). Note that the PSD (red) is adjacent to the protrusion, but not extending into it. Note also the close apposition of the pre- and postsynaptic cell membranes in the intruded region (inset in **a_1_**). Scale bar represents 0.5 µm. (**b**-**d**) 3D ssTEM reconstructed examples of synaptic boutons in which a cortical spine protrusion is invaginated into a labeled presynaptic Po bouton. A dashed white line highlights the approximate border between the spine protrusion and the remaining spine head. The synaptic bouton in (**b**) is a reconstruction made from the same series of sections shown in (**a_1_**-**a_4_**) (red lines). Scale bar represent 0.5 µm. (**e**) Bar histograms showing the distribution in PSD surface area in thalamocortical Po synaptic boutons in each of the three cortical regions examined. Total numbers of PSD are represented in pink. PSDs with a dendritic spine protrusion adjacent to them are highlighted in red. In the box, the total number of PSD measured in each cortical region, the percentage of PSDs with adjacent protrusions, and the mean surface area ± SD of the protrusion membrane are indicated. (**f**) To allow a direct comparison, PSD surface areas and protrusions measured in S1-L4 VPM synapses ^25^ are displayed in gray/black.

### Quantitative comparisons of structural synaptic parameters

To elucidate structural differences between Po axon synapses, parameters were quantitatively compared between the three cortical domains studied (Fig. 3d-h; Table 1). This analysis revealed striking differences. For example, Po boutons in MC-L4/3 were found to be significantly larger (∼60% in both volume and surface area) than Po boutons in S1-L5a (Fig. 3d). Most boutons in L1 were small, although the difference in average size was less apparent due to the presence of occasional large boutons interspersed in the axonal branches (Supplementary Fig. SM4). Mitochondrial volume per Po bouton was ∼33% larger in the MC-L4/3 than in the S1-L5a boutons, consistent with their bigger size (Fig. 3e). Most importantly, boutons in MC-L4/3 and S1-L1 contained vesicle pools about twice as large as those in S1-L5a (Fig. 3g). In addition, both the head volume and the PSD surface area of the spines postsynaptic to Po boutons in MC-L4/3 and S1-L1 were significantly larger (+50%) than those of spines postsynaptic to S1-L5a Po boutons (Fig. 3g-h).

Next, we compared the structure of synapses established by Po vs. VPM thalamic nuclei axons in S1 by putting side by side the 3D Po bouton measurements and those of VPM S1-L4 boutons ^25^. This comparison revealed that the Po-L5a boutons are −46% less in volume, and −33% in mitochondrial volume and contained −57% less synaptic vesicles than VPM-L4 boutons (Fig. 3d-f). In contrast, the Po MC-L4/3 boutons are statistically indistinguishable from VPM S1-L4 boutons with respect to these three parameters. Overall, this analysis demonstrates that a) axons from different thalamic nuclei can form structurally different presynaptic specializations in adjacent layers of the same cortical columns and b) axons from the same thalamic nucleus can form structurally different specializations in separate areas/layers.

In contrast, S1 spines postsynaptic to VPM (L4) and Po (L5a) boutons showed almost identical head volumes and PSD sizes. These two parameters were different, and overall much larger, for spines postsynaptic to S1-L1 (+83% and +45%, respectively) or MC-L4/3 (+50% and 54%) Po boutons (Fig. 3h), consistent with the notion that postsynaptic element differences reflect to a larger extent the local idiosyncrasies of distant neuropils.

Finally, to detect possible associations between pairs or groups of features in the structure of the Po boutons, we conducted a correlation (R^2^) and cluster analyses of the structural parameters analyzed. Synapses in non-varicose axonal zones were not included. These analyses revealed that in all cortical areas, particularly in MC L3/4, the volume of Po synaptic boutons correlates with that of their resident mitochondria, but much less so with the size of their vesicle pool (Supplementary Fig. SM5 and SM6). In each neuropil, pre- and postsynaptic parameters clustered independently; for example, the PSDs surface area and number were only weakly or not correlated with bouton size, but PSD surface area is positively correlated to the volume of the spine head (Supplementary Fig. SM5 and SM6).

### Varicosities in the MC branches of Po axons are consistently larger than those in branches of the same axons in S1

For the 3D electron microscopic study, relatively large populations of Po cell axons were anterogradely labeled with BDA. Since studies in rat indicate that MC and S1 may be targeted by axonal branches of the same Po neuron ^4, 24^, it remained unclear whether structural differences of Po synapses in MC and S1 reflect a) two different Po cell populations, each projecting either to MC or to S1, or b) area-specific synaptic structures in the collaterals of the same Po neuron. To address this question, isolated individual Po neurons were transfection-labeled with a Sindbis-pal-eGFP RNA construct using *in vivo* electroporation. From a larger collection of fully-reconstructed Po neurons projecting to a variety of cortical territories (n=12, data not shown), three cells were found to specifically innervate vibrissal motor cortex; remarkably, all three cells had, in addition, a collateral axon branch arborizing in the vibrissal region of the primary somatosensory cortex.

The arborization of two of these cell axons in S1 and MC are illustrated in Fig. 6; the third is shown in Supplementary Fig. SM7. The laminar distribution of these individual axons in the cortex was similar to that produced by bulk-labeling with BDA iontophoresis. Under light-microscopic analysis, MC-L4/3 boutons in the three neurons were consistently larger (1.13 ± 0.43 μm^2^, 1.13 ± 0.74 μm^2^ and 1.42 ± 0.49 μm^2^, respectively) than those formed by the branches of the same axons in S1-L5a (0.74 ± 0.30 μm^2^, 0.83 ± 0.39 μm^2^ and 0.96 ± 0.35 μm^2^) or S1-L1 (0.73 ± 0.37 μm^2^ and 0.91 ± 0.46 μm^2^), the third cell lacked axon branches in S1-L1 (see Fig. 6f-g and Supplementary Fig. SM7). Statistical comparison of mean bouton projection areas and size distributions revealed that the differences were highly significant (Fig. 6j-m). Hence, our analysis shows that the separate axonal branches of individual Po neurons form structurally different boutons in MC and S1.

**Fig. 6.**
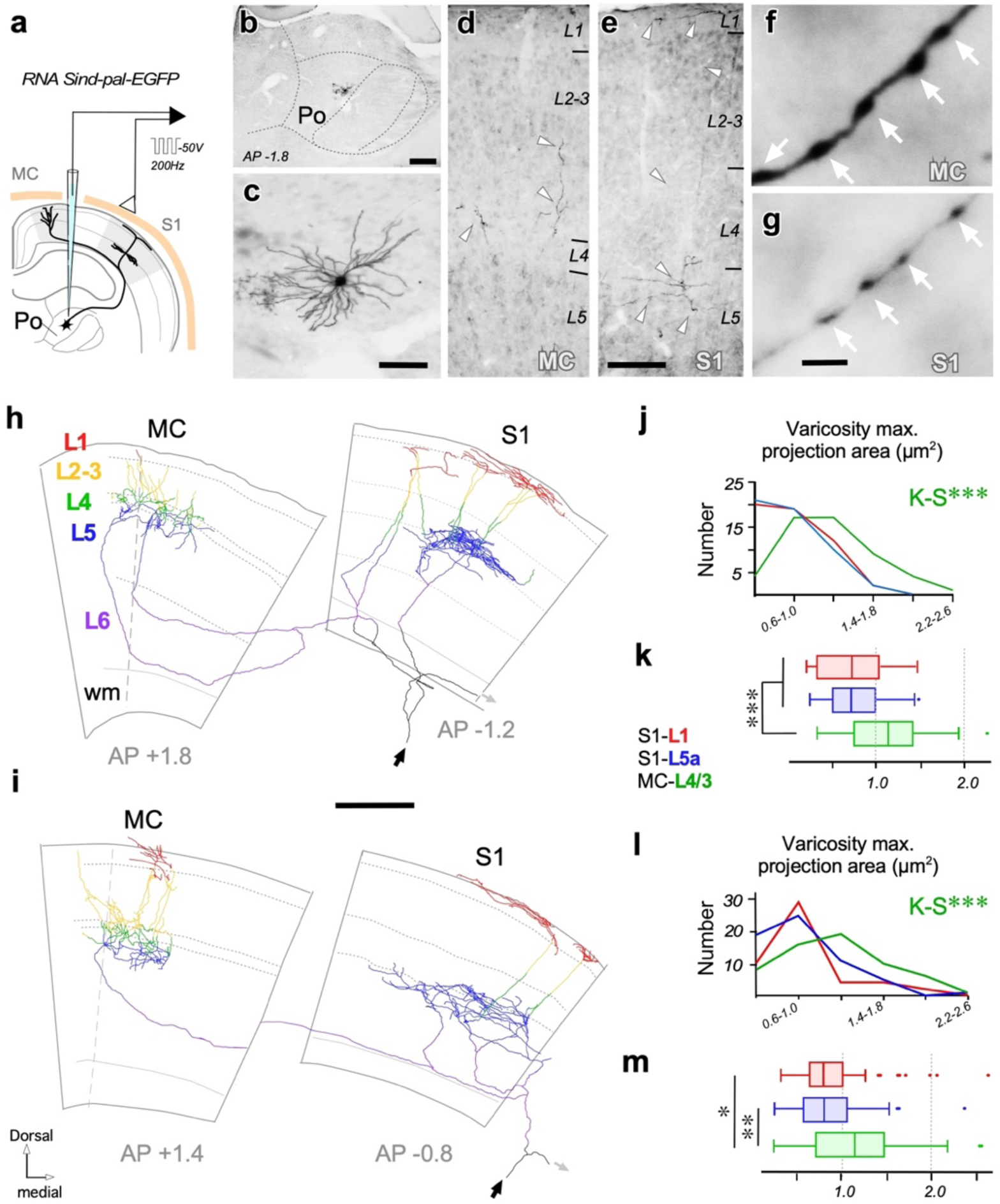
Individual branched Po axons target simultaneously MC and S1 and have varicosities of significantly different size in each area. (**a**) Schematic diagram of the Sindbis-pal-eGFP RNA electroporation procedure. (**b**) Coronal section through the thalamus showing an isolated neuron transfected in Po. (**c**) Somatodendritic morphology of the cell shown in “b”. (**d**, **e**) Labeled thalamocortical axon fragments (white arrowheads) as seen in columnar samples from MC (**d**) and S1 (**e**). (**f**, **g**) Po cell axonal boutons in MC or S1 at high light microscopic magnification. Note the marked differences in axonal diameter and varicosities sizes (white arrows). (**h**, **i**) Camera lucida reconstructions of the axonal arborizations of two individually-labeled Po neurons targeting both MC and S1. Axonal domains located in the different cortical layers or in the white matter (wm) are coded in different colors. Outlines of cortical layers (grey lines) are shown as a background. A dashed line in MC indicates border between areas Agl/M1 and Agm/M2 ^48^. Distance to bregma is indicated. The point of entry of the axon coming from the thalamus (black arrow) and a branch extending to more lateral areas (not shown; gray arrow) are indicated. (**j, l**) Comparison of varicosity size (maximal projection areas) distributions of the MC L4/3 vs. S1-L5a vs. S1-L1 branches of each axon. K-S = Kolmogorov-Smirnov. (**k, m**) Comparison of mean maximal projection area. Mann-Whitney U Test. Scales bars: b 250 µm; c 50 µm; d, e 100 µm; f, g 5 µm; h, i 500 µm.

## Discussion

Here, we demonstrate that individual Po neuron simultaneously innervate MC and S1 through branched axons that have varicosities (putative synaptic contacts) of markedly different size in each area. These structural differences were found to be even more pronounced using high-end 3D-electron microscopy: the Po boutons in MC-L4/3 differ significantly from Po boutons in S1 (L1 and L5a) in volume, surface area, and mean number of synaptic contacts. Moreover, MC-L4/3 and S1-L5a Po synapses are significantly different in vesicle pool size, in PSD surface area and shape, and in their proportion of synapses established on spines. Interestingly, about a quarter of the Po synapses occur in non-varicose axonal segments. Comparison with our previous data on VPM S1-L4 synapses ^25^ reveals both a sharp contrast between Po and VPM synapses in S1, which correlate with recently discovered differences in the functional properties of these synapses, as well as intriguing similarities between Po synapses in MC and VPM synapses in S1 (Table 1, Fig. 7).

**Fig. 7.**
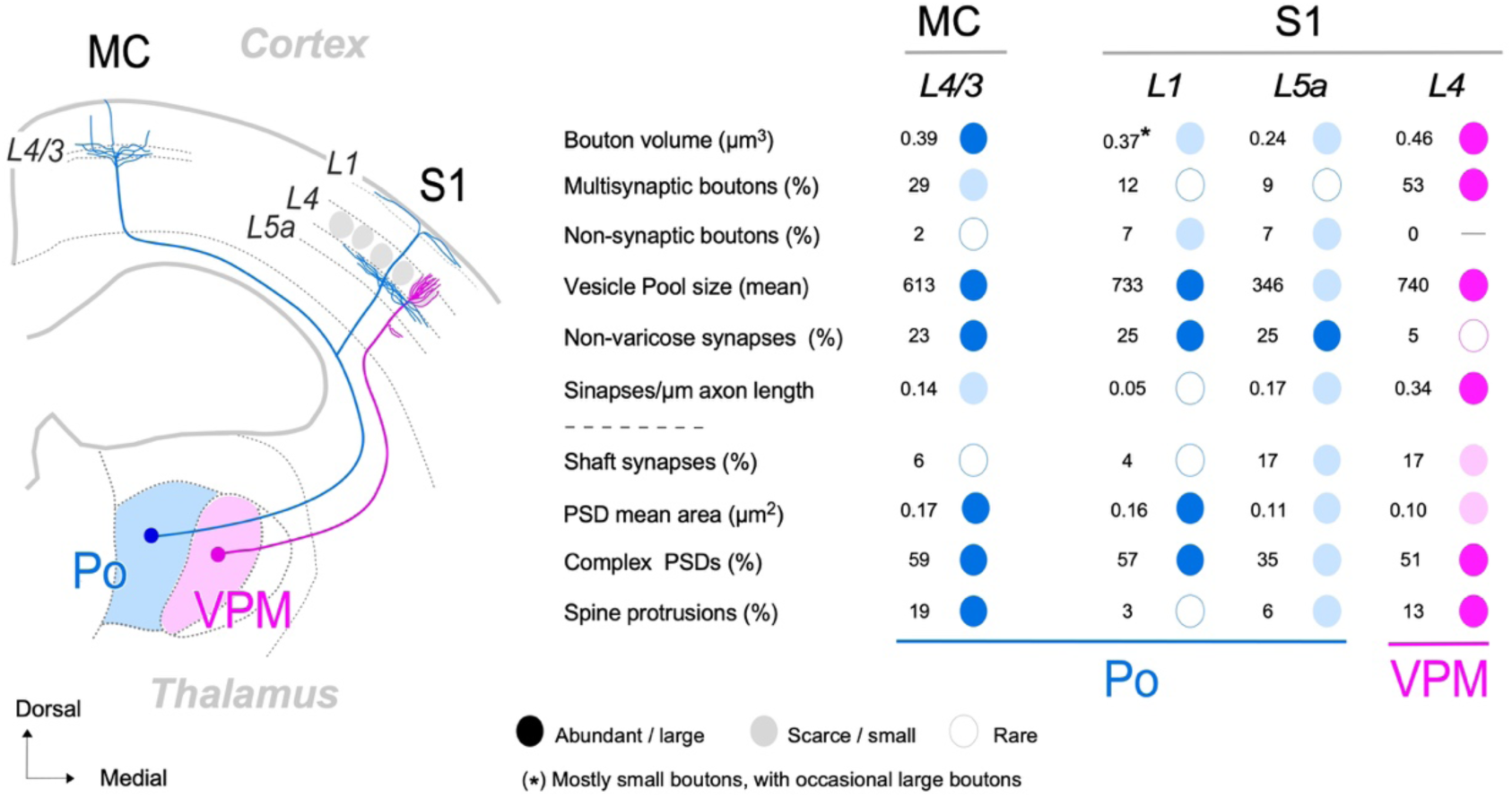
Graphic summary of relevant structural differences between Po axon synapses in MC-L4/3 and S1 (L1, and L5a) and comparison with the S1-L4 VPM axon synapses. The diagram on the left depicts, on a coronal mouse brain section, the trajectory and cortical arborizations from a Po cell axon (blue) and a VPM cell axon (magenta). The table on the right summarizes in simplified graphic format, to facilitate comparisons, the most salient differences in synaptic structure observed in the present study between Po boutons in MC-L4/3, S1-L1 and S1-L5a. For clarity, numbers are rounded-up mean values (see Table 1 for details). The column on the right (magenta) displays, with the same conventions, the values measured in S1-L4 VPM synapses.

### Individual Po axons form structurally different synapses in MC and S1

Thalamic inputs reach the cortex via a diverse array of overlapping axonal pathways. New techniques that make it possible to label the complete axonal arborization of individual neurons are revealing that axons from most thalamic nuclei branch to target several cortical areas (reviewed by ^31^). In rats, individual Po cell axons can innervate both the somatosensory and motor cortices, and often other regions ^4, 24^. In a recent study of anterogradely bulk-labeled mouse Po axons, we showed that axonal varicosities in MC-L4/3 are, as a population, significantly larger than varicosities in S1-L5a ^3^. Here, using single-cell transfection-labeling of single Po neurons and quantitative axonal and bouton 3D-volume reconstructions, we demonstrate that these large MC-L4/3 and small S1-5a varicosities occur on separate branches of individual Po axons, and dovetail much deeper differences in synaptic 3D ultrastructure.

The most striking differences between Po MC-L4/3 axon synapses and S1-L5a is the difference in their mitochondrial volume, PSD area and number of synaptic vesicles. Large bouton and mitochondria volume and large vesicle pools are all linked to the elevated energy supply and/or calcium homeostasis required to maintain high release rates ^28, 32, 33, 34^. Mitochondria boost local ATP generation and control Ca^+2^ levels which allow enhanced mobilization and recycling of synaptic vesicles for exocytosis, neurotransmitter release and for the generation of synaptic membrane potentials, specially under repetitive high-frequency firing ^33^. Likewise, the presence of large vesicle pools and extensive and complex PSDs is associated with higher neurotransmitter release probabilities and synaptic efficacy ^17, 18, 19, 35^ and to the number and distribution of postsynaptic receptors ^20, 22, 23^. The striking differences in these parameters between Po MC-L3/4 and S1-L5a boutons thus imply that the first may readily keep a high release probability at high-frequency firing rates, which is consistent with the observed capacity of MC Po synapses to transmit signals with higher efficacy and temporal acuity than Po S1 synapses ^3, 16^. Besides the structural differences, specific ionotropic and metabotropic receptor distributions may contribute to the temporal profile divergence of MC and S1 neuron responses to the repetitive activation of Po synapses ^3^.

Thalamocortical shaft synapses in rodent neocortex correspond in a large proportion to contacts on cortical inhibitory neurons ^36, 29^, but see ^37^. Thus, our data indicate that, mouse Po axons contact S1-L5a cortical interneurons in a proportion comparable to other thalamocortical systems and species, including VPM-L4 axons (Table 1, Fig. 7). However, we found three times less dendritic shaft synapses in MC-L4/3, suggesting that Po axons establish relatively fewer contacts on MC interneurons, a finding in register with the reported scarcity of VGluT2-possitive boutons contacting smooth dendritic shafts in mouse motor cortex ^29^. Thalamocortical axon contacts on cortical interneurons produce a powerful di-synaptic feedforward inhibition, in parallel with the excitation, on cortical neurons ^38, 39, 40^; a scarcity of Po shaft synapses in MC seem thus to be consistent with our recent observation that MC unit responses are facilitated by rapid repetitive Po axons activation, while the S1 responses become depressed ^3^.

### Po and VPM synapses in S1 are markedly different in structure

Comparison of Po axon S1 synapses with VPM axon S1 synapses (Fig. 7; see also ^25^) reveals a remarkable contrast: both types of thalamocortical axon contact postsynaptic shafts or spines in the same proportion, and their PSD sizes are similar. However, their presynaptic ultrastructure is strikingly different. Virtually all (95%) VPM synaptic boutons were large, containing one or several mitochondria, large vesicle pools, and most of them (53%) contained more than one (up to four) active zones. Only 5% of the VPM synapses lacked a mitochondrion (interbouton synapses), and no VPM axonal segment containing a mitochondrion inside lacked an active zone ^25, 27^ (Table 1; Fig. 7). In other words, in VPM axonal arbors, all mitochondria were found near a synapse. In contrast, Po boutons in S1-L5a were nearly 50% smaller, contained 30% smaller mitochondrial volumes and 50% smaller vesicle pools. Only 9% were multisynaptic, and these had, at most, two active zones. Moreover, about 25% of Po synapses (defined by a PSD and a vesicle pool) were found in “non-varicose” axonal domains. Remarkably, 7% of the varicosities contained a mitochondrion but lacked an active zone. In Po axons, therefore, mitochondria are far less concentrated around synaptic sites, a pattern reminiscent of that observed in the Shaffer cortico-cortical collateral system of the hippocampus, where over 50% of synaptic sites lack a mitochondrion and 8% of axonal varicosities containing a mitochondrion had no synaptic contact ^28^.

Po S1 and VPM L4 synapses evoke each markedly different temporal response profiles in cortical cells as a result of the selective involvement of different types of glutamate receptors: the VPM synapses involve only ionotropic receptors and their EPSCs depress markedly by repetitive stimulation ^9^, while Po synapses involve both ionotropic and metabotropic receptors and display EPSCs-facilitation ^3, 15^. Interestingly, the different presynaptic mitochondrial content in VPM vs. Po synapses might also contribute to their different effects on S1 neurons. By boosting local Ca^2+^ and ATP levels, mitochondria actively promote the docking and/or fusion of synaptic vesicles ^33, 41^, as well as the transport of vesicles from the resting to the recycling and readily releasable pool ^42^. As a result, mitochondria-rich synapses are capable of maintaining a high release probability over a wide range of firing rates ^34^. Our observations of larger mitochondrial concentration near the VPM S1 synapses seems thus consistent with electrophysiological evidence that VPM synapses produce relatively large (∼4.8mV) excitatory postsynaptic potentials ^9^ or currents ^10^ with low failure rates ^11, 12^, while Po synapses in S1-L5a produce smaller postsynaptic currents ^10, 15, 16^, that have slower rise and decay times ^14^.

In addition, studies over the past decade have revealed that mitochondrial distribution along axonal trees is optimized to match the local metabolic demands of their synapses. Importantly, a significant fraction of axonal mitochondria may remain mobile in adult axon arbors (reviewed by ^33^). Low ATP and high calcium levels promote the docking and/or fusion of mobile mitochondria near highly active synapses ^41^. From this perspective, the presence in S1 Po axons of many synapses without mitochondria and of mitochondria distant from synapses suggests that synaptic efficacy might be readily enhanced by the mobilization of mitochondria to particular synapses. In contrast, the pattern observed in in VPM S1 axons (high mitochondria concentration at synapses and absence of non-synaptic mitochondria) may not allow significant changes in efficacy though mitochondrial re-distribution. This scenario is consistent with the recently discovered capacity of Po S1-L5a synapses for delayed yet stable potentiation as a result of conditional adult learning, a capacity that is lacking in VPM S1-L4 synapses ^13^.

Finally, the comparison of the MC Po axon presynaptic structure of with that of VPM S1 synapses reveals intriguing similarities (Fig. 7; Table 1), despite the fact that these two synapse populations arise from different thalamic nuclei and are located in widely separated cortical domains. Such overall resemblance is consistent with electrophysiological observations that both Po MC and VPM S1 synapses elicit similarly large EPSCs ^9, 16^, exhibit paired-pulse depression and involve only ionotropic receptors ^3, 16^. Remarkably, however, the size of the Po MC-L3/4 synapse PSDs is 60% larger than that of the VPM L4 synapses, suggesting that the former may have even greater high synaptic strength and release probability. Moreover, as pointed out earlier, these synapses may have their effects on cortical cells less temporally curtailed by feedforward inhibition ^3^.

### Large non-synaptic spine intrusions are frequent in thalamocortical boutons

We show that many Po and VPM boutons have an elongated, thick protrusion of the postsynaptic spine head invaginated into them. Two previous 2D electron microscopic studies of lateral geniculate nucleus axon synapses in the primary visual cortex of ferrets ^43^ and tree shrews ^44^ reported similar spine profiles, suggesting that spine intrusions may be common in mammalian thalamocortical synapses. While their precise functional significance remains to be determined; our data already provide some intriguing clues. For example, invaginations are always found adjacent to the spine active zone, yet seem not to be directly related to its size (Fig. 5e-f). They are substantially more prevalent in the Po MC-L3/4 and the VPM S1-L4 synapses than in the S1 Po synapses (L5a and L1), suggesting some relationship with the absence of metabotropic glutamate receptors. Moreover, the large (up to 20 times the size of the active zone) and narrow invaginated intermembrane space is bound to create non-linear diffusion conditions, free of glial scavenging, for neurotransmitters, trophic factors or other secreted molecules. It is even possible that the extensive patch of parallel and closely apposed membranes might allow local electric field (ephaptic) conduction ^45^ between the cortical spine and its thalamocortical bouton.

In conclusion, we have demonstrated here that differences in the composition of synaptic structure underlie and explain the divergent responsivity and plasticity of MC vs. S1 neurons to Po input ^3, 16^, as well as that of S1 neurons to VPM vs. Po inputs ^10, 13^. Moreover, the evidence that these structural and functional differences actually occur between synapses in separate branches of the same individual Po cell axons indicates that current global models of thalamo-cortical wiring and interaction ^46, 47^ may require revision.

## Acknowledgements

The authors gratefully acknowledge, Ms. Begoña Rodriguez and Ms. Marta Callejo for excellent technical help.

## Author Contributions

JRM, FC, JHRL and CP designed the experiments,

JRM, CP and MRT prepared the materials,

JHRL, JDF, AM, AS and LAN provided or operated key equipment,

JRM, CP, MRT, DCT, AS, RY and AR analyzed the data,

JRM, FC, CP and RY prepared the figures and tables,

FC, JRM, and JHRL wrote the paper with input from DCT, AR, AM and JDF.

## Competing Interests statement

This work was supported by funding from the European Union’s Horizon 2020 Research and Innovation Programme (Grant Agreement No.785907 HBP SGA2). Additional support was provided by Spain’s Ministerio de Ciencia, Innovación y Universidades (BFU 2107-88549-P) to FC, and by grants of the Helmholtz Society (JHRL).

All the authors declare that they have no other financial interests that might be relevant to the submitted study.

The data that support the findings of this study are available from the corresponding author upon reasonable request. In addition, all serial electron microscope image stacks and the two short videos cited in Supplemantary Materials Figure SM3 will be made available before publication in the Human Brain project Graph Data Platform https://www.humanbrainproject.eu/explore-the-brain/search” under a CC-BY license.

## Methods

### Animals & anesthetic procedures

Experiments were performed on adult (60-105 days old, 25-32 g in body weight) male C57BL/6 mice bred in the Autónoma de Madrid University colony. All procedures involving live animals were conducted under protocols approved by the University ethics committee and the competent Spanish Government agency (PROEX175/16), in accordance with the European Community Council Directive 2010/63/UE. Animals were housed in pairs in cages containing some toys, and provided chow and water ad libitum under a 12 hours light/dark cycle. Efforts were made to minimize the number of animals required. Six male mice (60-65 days in age) were used for the experiments aimed at BDA population-labeling of Po axons for light- and electron microscopy. Thirty-two further mice were electroporated with Sindbis Pal-eGFP RNA to reveal the full extent of individual Po cell axons.

For both types of experiment, mice were first anesthetized with an initial intraperitoneal injection of ketamine (0.075 mg/g body weight) and xylazine (0.02 mg/g body weight), and then maintained with oxygenated isoflurane (0.5-2%) throughout the surgical procedure. Animals recovered promptly after the interruption of the isofluorane flow, at the end of the surgical procedure. Ibuprofen (120 mg/l) was added to the drinking water to ensure analgesia during the survival period. At the time of sacrifice, animals were overdosed with sodium pentobarbital (0.09 mg/g body weight, i.p.).

### BDA iontophoresis for selective population-labeling of Po axons

Animals were placed in a stereotaxic frame (Kopf Instruments, Tujunga, CA, USA). BDA10K (2.5% in saline; Molecular Probes-Invitrogen, Carlsbad, CA, USA) was iontophoretically microinjected (positive tip, 0.7–0.8 µA; 1 sec on/off cycle, 30–40 min duration) with a Midgard Precision Current Source (Stoelting, Wood Dale, IL, USA) under stereotaxic guidance (1.8 mm posterior, 1.3 mm lateral, and 2.7 mm ventral to Bregma ^48^), using borosilicate micropipettes (WPI, Sarasota, FL, USA; outer tip diameter: 7–10 µm). The micropipette was finally removed, and the muscle and skin were sutured and disinfected with iodinated povidone.

### Tissue processing of the BDA-injected brains

Following a 5-days survival, injected mice were perfused with phosphate-buffered saline (0.1 M PBS) followed by a fixative of 4% paraformaldehyde and 0.1% glutaraldehyde in 0.1 M PB for 30 min. After the perfusion, brains were removed from the skull, and post-fixed by immersion for 1 hour at 4°C in the same fixative. Two series of parallel coronal sections (50 µm-thicknesses) were cut with a Leica VT 1200S vibratome (Leica Microsystems, Nussloch, Germany). Sections were then cryo-protected by incubation in a sucrose solution (30% in 0.1M PB) overnight, and were then rapidly freeze-thawed in liquid nitrogen (1 min) and stored in PB until further use.

In the first series of sections, peroxidase activity was blocked by incubation in PB-buffered H_2_O_2_ for 10 min, and sections were then incubated in an avidin-biotin-peroxidase kit (1:100; Vectastain Elite^TM^, Vector Laboratories, Burlingame, CA, USA) diluted in PB. After washing in PB, labeling was visualized using the glucose oxidase-3-3’diaminobenzidine (DAB) nickel sulfate-enhanced method ^49^. Sections were mounted onto gelatin-coated glass slides, counterstained with thionin, dehydrated in graded ethanol, defatted in xylene, and coverslipped with DePeX (Serva, Germany). These sections were used for light microscopic analysis (Nikon Eclipse 600, 4-40X objectives) of the cytoarchitectonic localization of the injection site in the thalamus and the anterogradely labeled axons in the neocortex. Out of the twelve BDA injected hemispheres, four showed BDA injections restricted to Po (Fig. 1A) and robust axonal labeling in S1 and MC (Fig. 1, Supplementary Fig. SM1). The two cortical areas and respective target layers were identified based on the thionin counterstaining. The vibrissal region of the MC as defined in this study is located along the border between cytoarchitectonic areas AgM/M2 and AgL/M1 of the frontal cortex ^3, 5^ (Bregma AP: +0.5 - +2.2 mm, ML: 1-1.5 mm). The vibrissal domain of S1 is readily identifiable by its barrels in L4.

The second series of sections followed a protocol identical to that described above, except for the omission of the oxygen-peroxide blocking step and the absence of the nickel sulfate-enhancement in the glucose-oxidase-DAB reaction. In the four injection experiments that produced optimal labeling of Po axons (see above), free-floating sections from this series containing the regions with the cortical labeling underwent further processing for EM. In these sections, the glucose oxidase-DAB reaction was followed by incubation in 1% osmium tetroxide (Electron Microscopy Science, Hartfield, PA, USA) diluted in PB for 45 min at room temperature. Following thorough washing in PB, sections were first rinsed in 50% ethanol, incubated for 40 min in 1% uranyl-acetate diluted in 70% ethanol in the dark, and dehydrated in an ascending series of ethanol to absolute ethanol. The dehydrated sections were transferred to acetonitrile (Scharlab, Barcelona, Spain), and then transferred to an epoxy resin (Durcupan^TM^, Electron Microscopy Science, Hartfield PA, USA) overnight. Finally, sections were flat-embedded in Durcupan^TM^ and polymerized at 60°C for 48 hrs.

After light microscopic inspection, samples containing dense labeling in the S1 and MC neocortex were cut out and glued onto pre-polymerized resin blocks for serial section and subsequent ssTEM imaging (Supplementary Fig. SM2).

### Tissue processing for ssTEM imaging of BDA-labeled Po boutons

Embedded tissue blocks that contained L5a and L1 in S1 barrel cortex or L4/3 in MC were cut with a Leica Ultracut UCT ultramicrotome (Leica Microsystems, Nussloch, Germany) into serial 60 nm ultrathin sections (around 90 sections/series). They were collected on pioloform-coated single-slot copper grids (Electron Microscopy Science). Thereafter, they were treated with lead citrate (3 min), and examined with a Libra 120 transmission electron microscope (EM, Fa. Carl Zeiss, Oberkochen, Germany) equipped with a Proscan 2K bottom-mounted digital camera (Fa. Albert Tröndle, Moorenweis, Germany) and the SIS Multi Image Acquisition software package (Olympus, Hamburg, Germany). At the EM level, BDA-labeled axons were easily identifiable by the opaque DAB reaction product. Serial digital images were taken at a magnification of 8000x and stored in a database until further use.

### FIB-SEM 3D tissue preparation and imaging of BDA-labeled Po boutons

In addition, some Durcupan-embedded tissue blocks from the same experiments were used to obtained 3D tissue samples using combined focused gallium ion beam milling and scanning electron microscopy (FIB-SEM). Here, a Crossbeam Neon40 EsB electron microscope equipped with a gallium FIB and a high-resolution scanning emission SEM column (Carl Zeiss) was used. In order to accurately select the regions of interest, a secondary electron microscopic image was acquired from the block surface that was overlaid and collated with previously obtained light microscopic images ^50^. Once the appropriate location was chosen, a gallium ion beam was used to mill the sample to allow visualization of brain tissue under the block face on a nanometer scale. The recently milled surface was then imaged using a back-scattered electron detector (1.8 kV acceleration potential). The milling and imaging processes were sequentially repeated in an automated way, providing a stack of serial digital images that represented a 3D sample of the tissue ^51^. Image resolution in the xy-plane was 5 nm per pixel. Resolution in the z-axis (section thickness) was 20 nm, so the voxel size of the resulting image stack was 5×5×20 nm.

With the above resolution parameters, images of 2048 x 1536 pixels (field of view of 10.24 x 7.68 microns) were obtained. A total of twenty different stacks of images of the neuropil in the three cortical regions of interest were obtained (seven stacks from S1-L5a, six from S1-L1, and seven from MC-L3/4). The number of serial sections per stack ranged from 75 to 478; the total number of serial sections was 4,874 (mean: 243.7 sections per stack). Registration (alignment) of serial sections was performed with the freely available Fiji software ^52^, using a rigid body model that allowed no deformation of individual images.

### 3D-volume reconstruction and analysis of ssTEM and FIB-SEM image stacks

The 3D-volume reconstructions and measurements on ssTEM images were carried out with the OpenCAR software (Contour Alignment Reconstruction) ^53^. Digital images were aligned creating an image stack where all structures of interest were defined by closed contour lines of different color.

The Z-stack image series acquired with FIB-SEM were 3D-segmented and measured with the Espina Interactive Neuron Analyzer software (v.2.1.10; freely available at http://cajalbbp.es/espina/) ^54^. 3D-reconstructions generated from separate Z-stacks were digitally stitched using Unity 3D modeling software (Unity Technologies, San Francisco, CA, USA).

The geometry of Po synaptic boutons, the PSD, mitochondria and the total pool of synaptic vesicles within individual boutons, and the postsynaptic target structures contacted by the synaptic boutons were completely 3D-reconstructed. From the contours, 3D-volume reconstructions were performed, from which surface area and volume measurements were obtained. The number of synaptic vesicles was estimated by a Physical Dissector stereological method ^55^ in Po presynaptic boutons with a light DAB reaction product.

For each structural parameter, a mean ± standard deviation (SD), the median, maximum and minimum values and a coefficient of variation (CV) were calculated. For subsequent data analysis for multiple comparisons, we used one-way analysis of variance (ANOVA) plus T3 Dunnet’s as a *post hoc* test with SPSS software (version 24; IBM, Armonk, New York). The threshold level of significance was set at *P < 0.05*, indicating this as (*), *P < 0.01* as (**) and *P < 0.001* as (***).

### Cluster analysis comparison of synaptic bouton ultrastructural parameters

All synaptic boutons reconstructed were clustered according to six structural parameters investigated, namely: bouton volume, bouton surface area, number of PSDs/bouton, number of mitochondria/bouton, volume of mitochondria and PSD surface area. The cluster analysis was performed using MATLAB and Statistics Toolbox Release 2016b (The MathWorks, Inc., Natick, MA, USA). The aim was to identify the synaptic parameters that best characterized the synaptic boutons investigated. Thus, a principal component analysis (PCA) was performed to simplify the dataset and to convert a set of observations of possibly correlated variables into a set of values of linearly uncorrelated variables called principal components (PCs) ^56^. Then, we performed a hierarchical cluster analysis (HCA) ^57^ on the new simplified dataset composed of the PCs. This method is used for unsupervised machine learning when the original data were unlabeled (For details see ^58^).

### Sindbis-pal-eGFP RNA electroporation for single-cell labeling

To directly visualize the complete axonal tree of individual mouse Po cells in an unambiguous manner, isolated Po neurons were transfected by means of *in vivo* electroporation ^59^ of a RNA construct engineered to drive the expression of an enhanced green fluorescent protein (eGFP) fused with a palmytoilation motif from the growth-associated protein 43 (GAP43) under the Sindbis viral subgenomic promoter (Sind-Pal-eGFP) ^60^.

Micropipettes were pulled from Kwick-Fill borosilicate capillaries (1 mm outer diameter; WPI). Inner tip diameter was adjusted to 10–15 µm. To eliminate RNAse activity, micropipettes were then kept in a stove overnight at 240°C and, after cooling, were backfilled with a RNA stock solution (1.8–2 µg/µl in 0.5M NaCl) and mounted on a holder (WPI) that has both a pressure port and electrode connection. All procedures were performed over clean, single-use surfaces, and metal instrument tips were briefly exposed to a flame.

The micropipette tip was stereotaxically positioned into Po. 50–100 nl of solution were injected using a precision electro-valve system (Picospritzer II, Parker Hannifin, Cleveland OH). Two to four 200 Hz trains of 1 ms negative-square pulses at 50 V were then applied through the micropipette tip using a CS20 stimulator (Cibertec, Madrid, Spain). The micropipette tip was left in place for 5 min before removing it from the brain. Finally, the bone defect was closed, the scalp sutured, and animals were returned to their cages.

Following 58–60 h survival, animals were overdosed with pentobarbital (80 mg/kg body weight, i.p), and perfused with saline (1 min), followed by 4% paraformaldehyde in 0.1 M PB, pH 7.4, for 8 min. Their brains were then removed, immersed overnight in the same fixative at 4 °C and cryoprotected by soaking in a sucrose solution (30% in PB) for further 24 hrs. Serial 50-µm-thick coronal sections were cut on freezing microtome.

To allow intensive high-magnification microscopic analysis, labeling was made stable and opaque by using immunohistochemistry against eGFP and glucose oxidase-nickel enhancement ^49^. Free-floating sections were incubated in rabbit anti-GFP serum (1:500; EXBIO, Prague, Czech Republic), followed by incubation with a biotinylated goat anti-rabbit serum (1:100; Sigma-Aldrich, St. Louis, MO, USA) and an avidin-biotin-peroxidase kit (1:100; Vectastain Elite, Vector Laboratories). Sections were then serially mounted onto gelatin-coated glass slides, air-dried, lightly counterstained with thionin, dehydrated and coverslipped with DePeX (Serva).

Sections were examined under brightfield optics at 10-40X. Labeled axons of Po neurons were found to span >30 serial coronal sections. Complete 2D-reconstructions of each axon were drawn using a Nikon camera lucida, scanned, and redrawn using a vector graphics software (Canvas X, ACD Systems, Saanichton BC, Canada).

Under bright-field optics, axonal branches appeared as sharply labeled filaments with frequent varicose swellings. To estimate and compare the size of varicosities (putative synaptic boutons), the maximal projection area was measured from live images using a Nikon DMX1200 camera fitted to the microscope and the NIS-Elements imaging software tools (v3.2; Nikon). To this end, the major perimeter of each varicosity was focused at 1000X and delineated over the computer screen using the “polyline” and “polygon” software tools. For each cortical area, layer and axon, 50 randomly selected varicosities were measured. Varicosities with cross-sectional areas (maximal projection) near or below the microscope resolution limit (< 0.2 µm^2^) we not included. Statistical analysis was performed with the SPSS software (IBM). Comparison of mean sizes of varicosities between areas/layers was performed using Mann-Whitney U test. Two-sample Kolmogorov-Smirnov test (K-S) was used to compare size distributions of varicosities between areas/layers.

## Supplementary Information

**Supplementary Fig. SM1.**
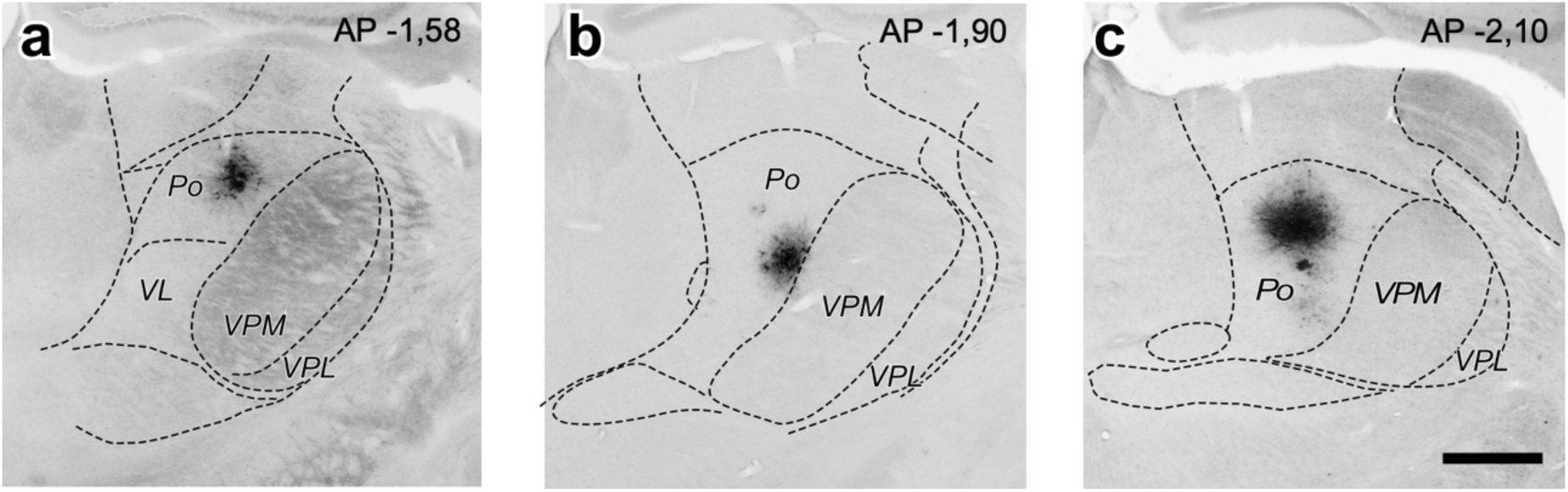
Three further examples of BDA iontophoretic deposits used to selectively label Po thalamocortical axons in MC and S1. (**a**-**c**) Coronal thalamus sections show the center of the BDA deposit in Po. Light thionin counterstain. Distance (in mm) caudal to bregma is indicated in the upper right corner. Scale bar represents 250 µm.

**Supplementary Fig. SM2.**
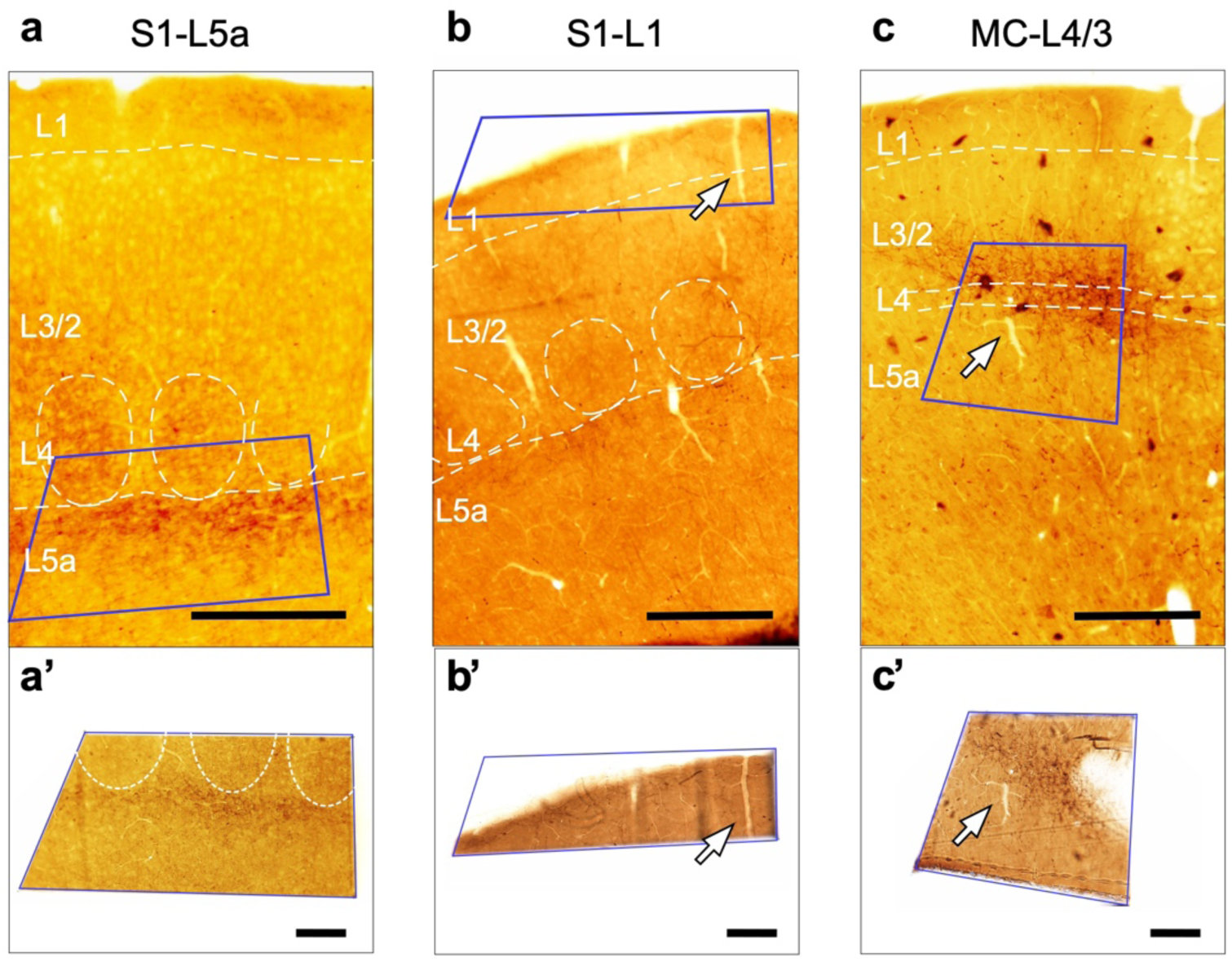
Selective sampling of tissue areas containing BDA-labeled Po axon branches. (**a**-**c**) Resin-embedded coronal sections. The blue rectangle indicates the area that was cut out for subsequent EM serial sectioning. Scale bars represent 250 µm (**a’**-**c’**). An arrow identifies the same vascular landmark in both images Scale bar represents 100 µm.

**Supplementary Fig. SM3.**
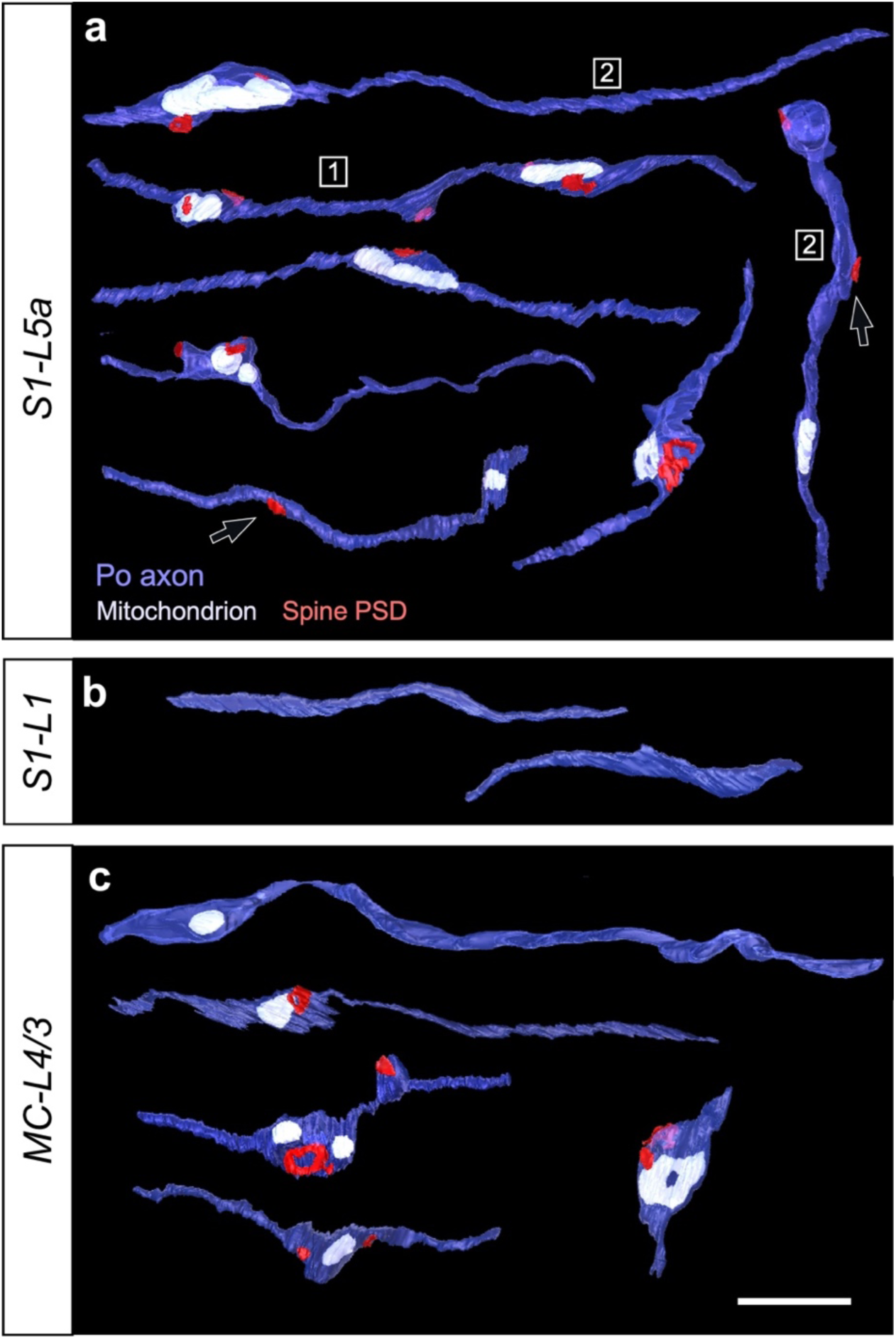
Additional thalamocortical Po axonal segments 3D-reconstructed from FIB-SEM image stacks. (**a**-**c**) Individual axonal segments from (**a**) S1-L5a, (**b**) S1-L1, or (**c**) MC-L4/3. All the PSDs in these particular segments were located on spines. Black arrows highlight PSDs located in non-varicose axonal domains. Axon segments labeled with numbers 1 and 2 are shown, within their correspondent 3D FIBSEM image stacks, in Supplementary videos SMV1 and SMV2. Scale bar represents 2 µm.

**Supplementary Fig. SM4.**
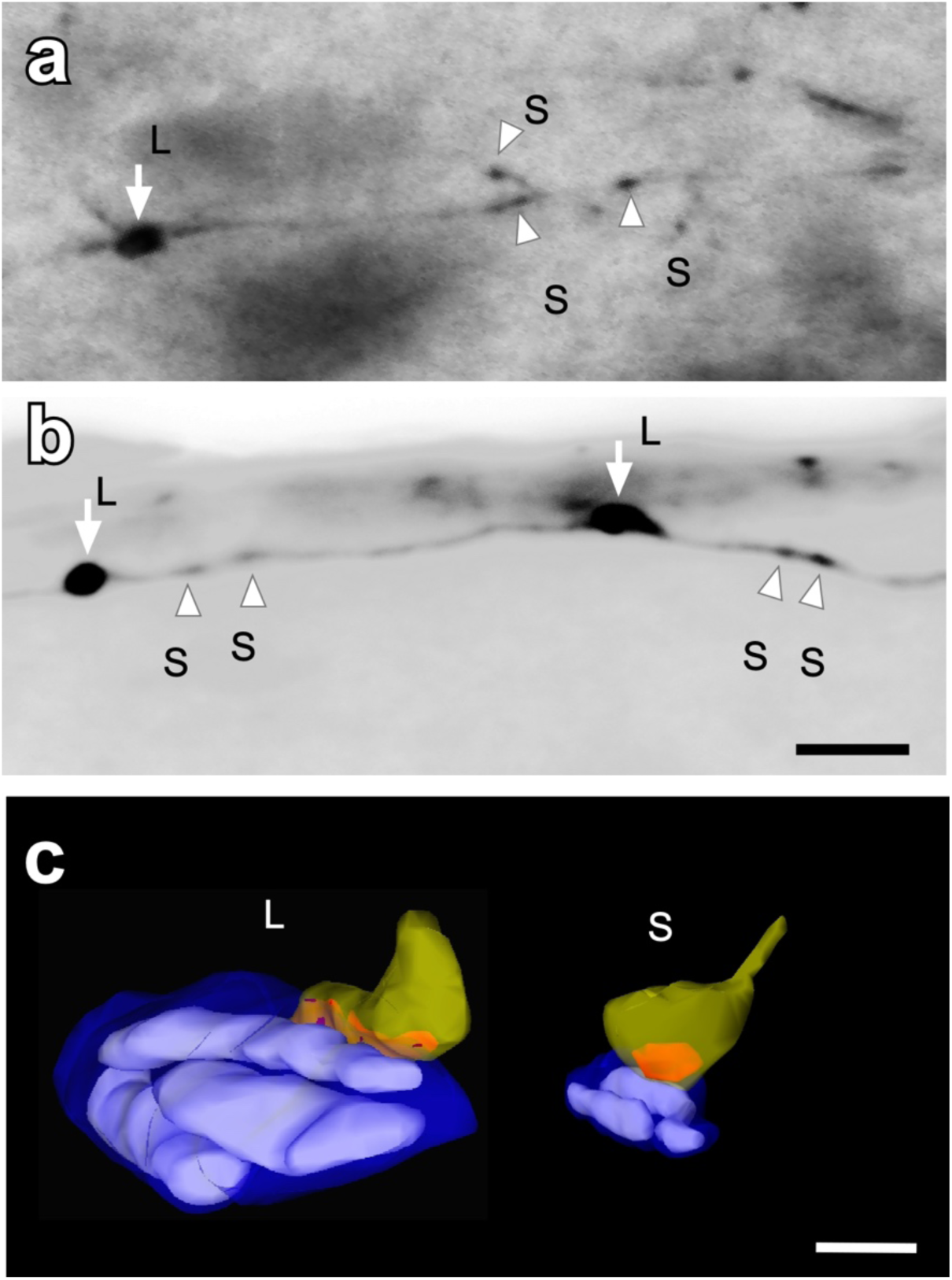
Po axon branches in S1bf L1 contain a majority of small synaptic boutons with occasional large boutons interspersed. (**a-b**) Light microscopy high-magnification images of a labeled Po axon containing both large bouton and small boutons. The branch in (a) was labeled by a BDA injection in Po. The branch in (b) was labeled by the transfection of an isolated Po neuron with Sindbis-pal-eGFP RNA. Scale bar represents 0.5 µm. **(c)** Representative examples of ssTEM 3D reconstructions of a larger (L) and smaller (S) Po axon boutons in S1-L1. Scale bar represents 2 µm.

**Supplementary Fig. SM5.**
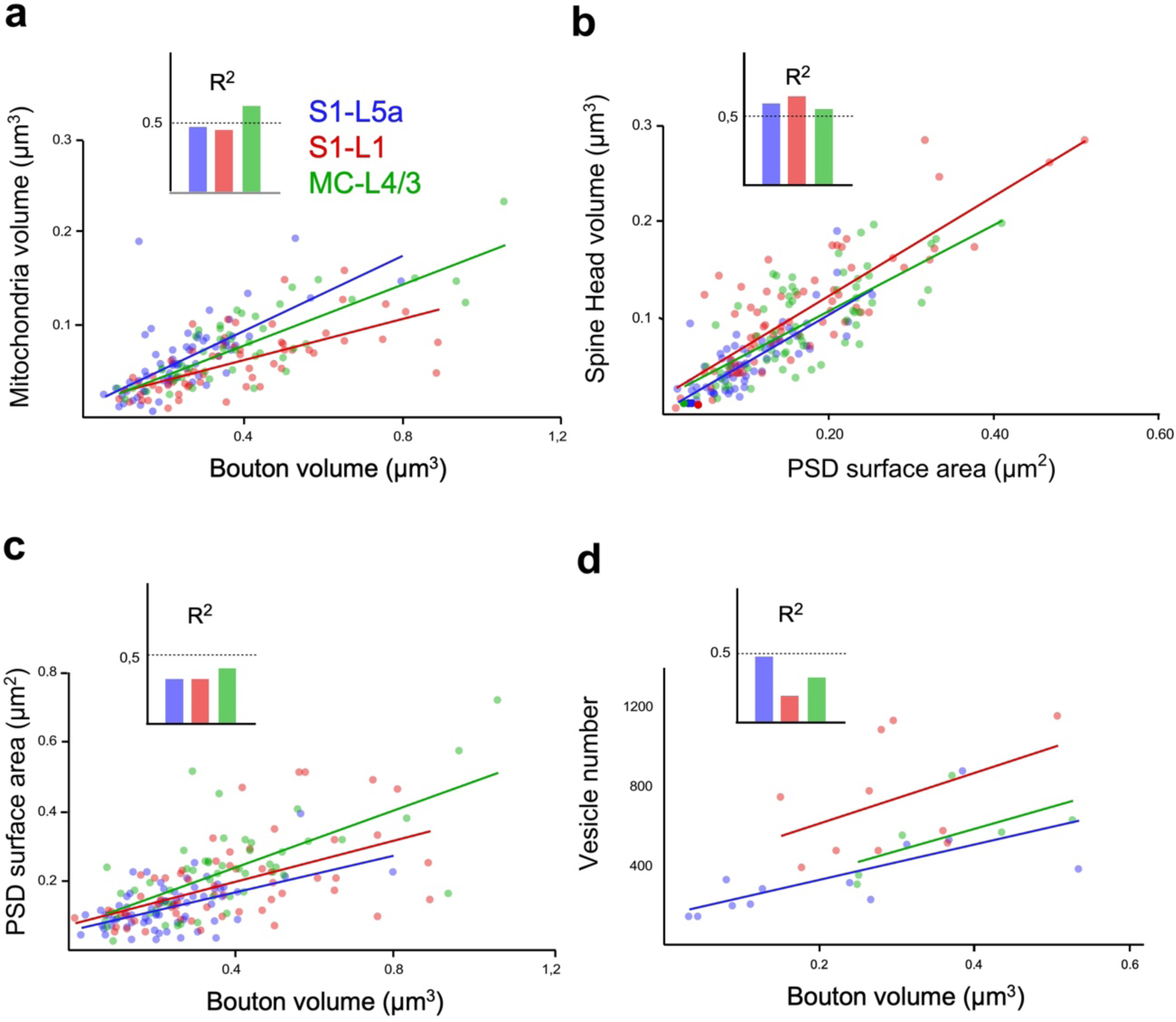
Correlation analyses between structural parameters of Po synaptic boutons in the three neuropil regions analyzed. (**a**-**d**) Dot plots showing the correlations between: (**a**) bouton volume vs. mitochondrial volume; (**b**) PSD surface area vs. spine head volume; (**c**) bouton volume vs. PSD surface area; and (**d**) bouton volume vs. number of synaptic vesicles. Correlations are indicated by linear regression as well as by the coefficient of determination (R^2^, bar histograms).

**Supplementary Fig. SM6.**
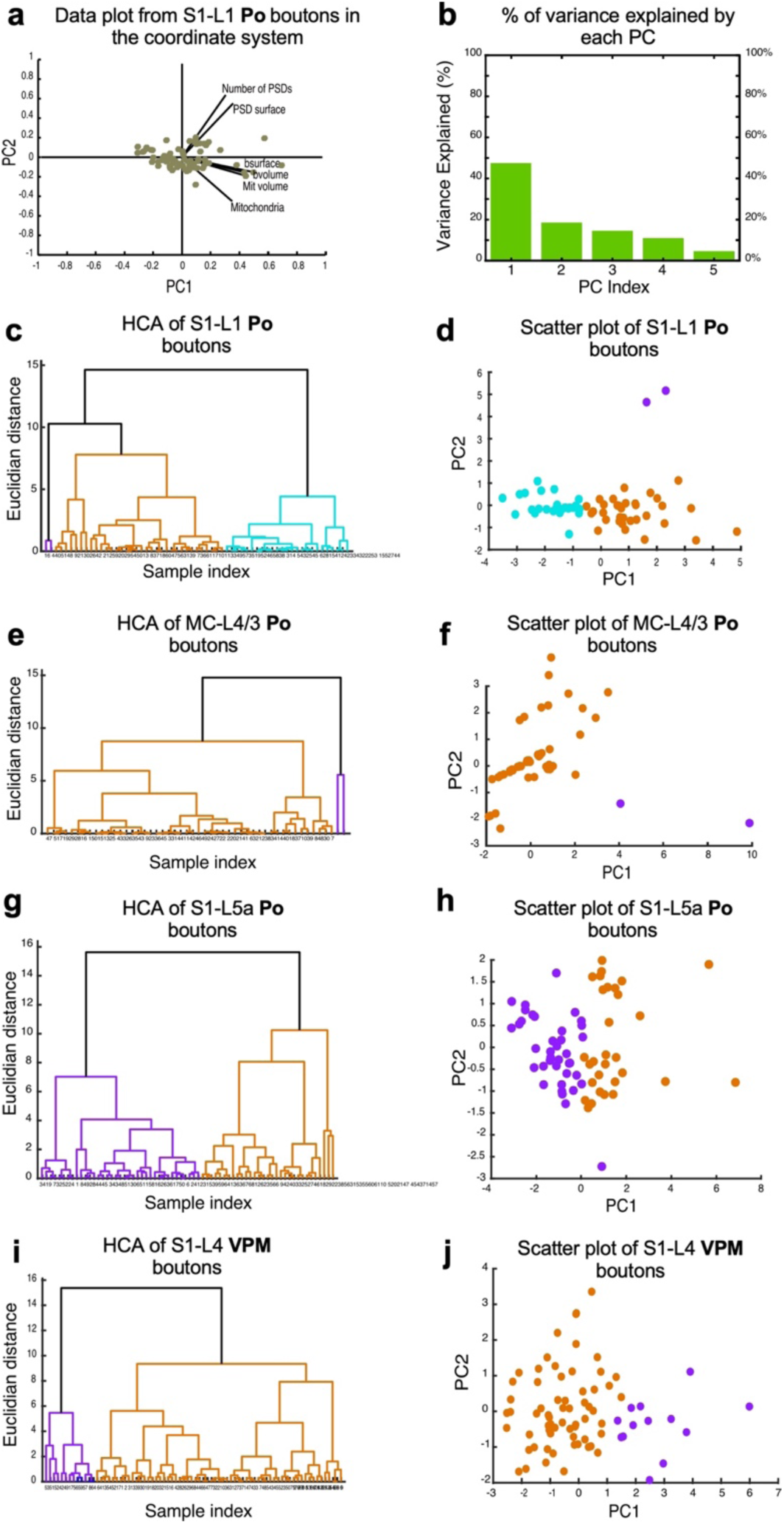
Cluster Analysis. (**a**) Projection of the original data of S1-L1 Po synaptic boutons in the new coordinate system based on the Principal Components (PCs), showing the coefficients for each variable and scores for each observation. Abbreviations used: *bvolume*: bouton volume; *bsurface*: bouton surface area; *Mit volume*: mitochondrial volume. (**b**) Histogram of the percentage of variance explained by the PCs for data from S1-L1 Po synaptic boutons. The PCs considered as main PCs are 1 and 2 as their total explain variance is about 70%. (**c**-**j**) Dendrogram and scatter plot analyses of Po synaptic bouton parameters in each of the cortical regions examined. (**c**, **d**) Data plots for S1-L1; (**e**, **f**) Data plots for MC-L4/3; (**g**, **h**) Data plots for S1-L5a. In addition, the same type of data plots for VPM boutons in S1-L4 ^25^ is shown in panels (**i**, **j**). In all plots, orange and purple colors are used to show the major clusters with respect to the synaptic parameters analyzed. Dissimilarity between clusters is indicated by the Euclidean distance.

**Supplementary Fig. SM7.**
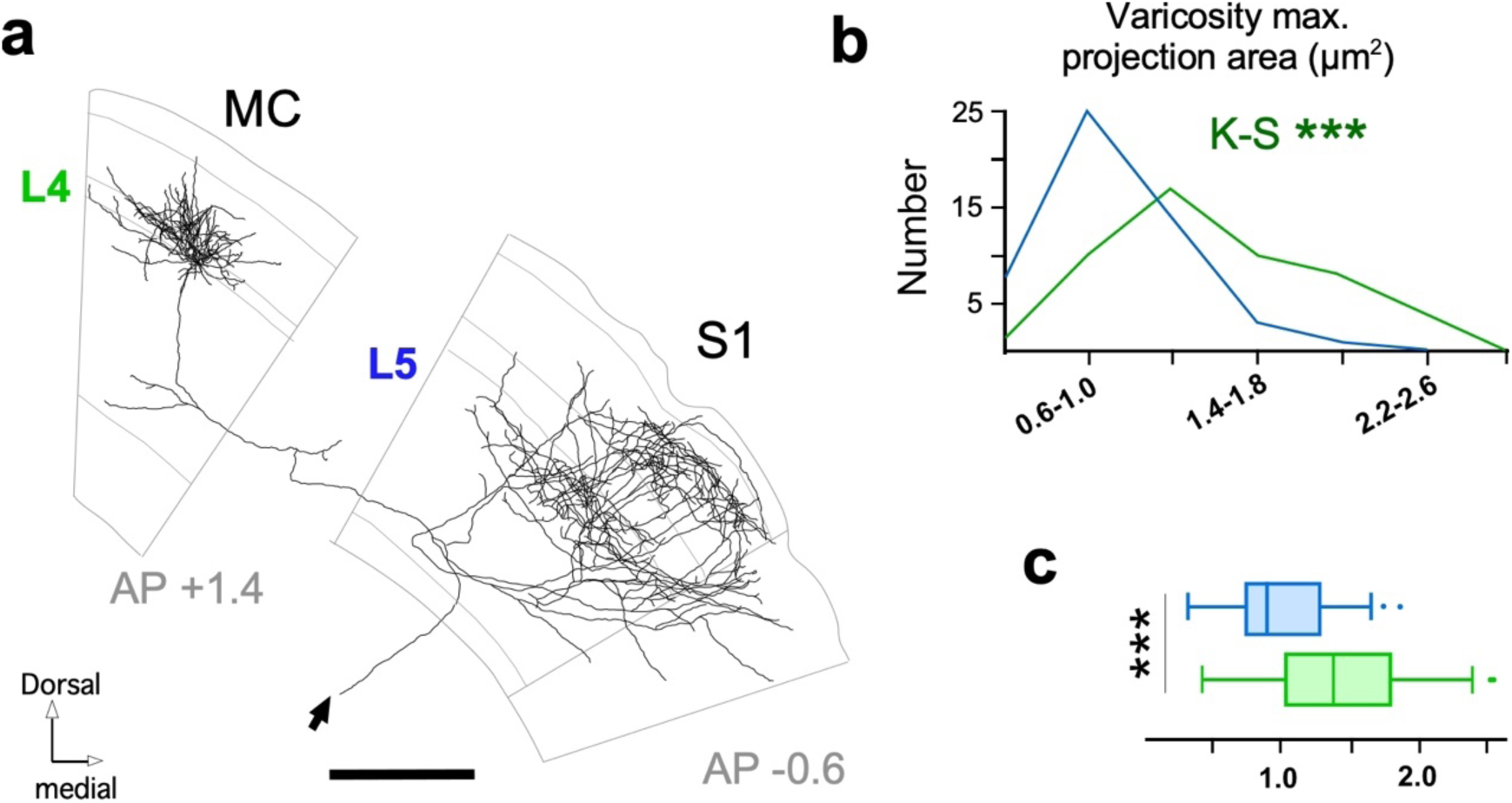
Axon reconstruction and varicosity size analysis of a further transfection-labeled Po neuron projecting both to MC and S1. (**a**) Camera lucida reconstruction of the axonal arborizations and projection pattern of an individually-labeled Po neuron targeting both MC and S1. For anatomical reference, outline sketches of cortical layers (grey lines) are shown as a background and the approximate distance to bregma (rostral “+” or caudal “-“, in mm) is indicated. The black arrow indicates the point of entry in the cortical hemisphere of the axon coming from the thalamus. Scale bar represents 500 µm (**b**) Comparison of varicosity sizes distributions (maximal projection areas) of the MC L4/3 vs. S1-L5a branches of the Po axon. This particular cell had virtually no axon branches in S1-L1. K-S = Kolmogorov-Smirnov; (**c**) Comparison of mean maximal projection area. Mann-Whitney U Test. (*): p <0.05; (**): p <0.01; (***): p< 0.001.

**Supplementary video SMV1.** Video showing an example of thalamocortical Po axonal segments 3D-reconstructed in S1-L5a from FIB-SEM image stacks.

**Supplementary video SMV2.** Video showing an example of thalamocortical Po axonal segments 3D-reconstructed in S1-L5a from FIB-SEM image stacks.

These videos will be available will be made available before publication in the Human Brain project Graph Data Platform https://www.humanbrainproject.eu/explore-the-brain/search” under a CC-BY license.

